# Discovery of a Novel Chemotype as DYRK1A Inhibitors against Alzheimer’s disease: Computational Modeling and Biological Evaluation

**DOI:** 10.1101/2023.11.03.565431

**Authors:** Nianzhuang Qiu, Chenliang Qian, Tingting Guo, Yaling Wang, Hongwei Jin, Mingli Yao, Mei Li, Tianyang Guo, Yuli Lv, Xinxin Si, Song Wu, Hao Wang, Xuehui Zhang, Jie Xia

**Affiliations:** School of Pharmaceutical Sciences, Shandong First Medical University& Shandong Academy of Medical Sciences, Taian, Shandong 271016, China; State Key Laboratory of Bioactive Substance and Function of Natural Medicines, Institute of Materia Medica, Chinese Academy of Medical Sciences and Peking Union Medical College, Beijing 100050, China; School of Pharmacy, Jiangsu Ocean University, Lianyungang, Jiangsu 222005, China; Beijing Tide Pharmaceutical Co., Ltd, Beijing 100176, China; State Key Laboratory of Natural and Biomimetic Drugs, School of Pharmaceutical Sciences, Peking University, Beijing, 100191, China

**Keywords:** Virtual screening, Molecular docking, DYRK1A, Alzheimer’s disease, tau hyperphosphorylation, Aβ aggregation

## Abstract

Dual specificity tyrosine phosphorylation-regulated kinase 1A (DYRK1A) plays an essential role in tau and Aβ pathology closely related to Alzheimer’s disease (AD). Accumulative evidence has demonstrated DYRK1A inhibition is able to reduce the pathological features of AD. Nevertheless, there is no approved DYRK1A inhibitors for clinical use as anti-AD drugs. This is somewhat the lack of effective and safe chemotypes of DYRK1A inhibitors. To address this issue, we carried out *in silico* screening, *in vitro* assays and *in vivo* efficacy evaluation with the aim to discover a new class of DYRK1A inhibitors for potential treatment of AD. By *in silico* screening, we selected and purchased 16 potential DYRK1A inhibitors from the Specs chemical library. Among them, compound **Q17** (Specs ID: AO-476/40829177) potently inhibited DYRK1A. The hydrogen bonds between compound **Q17** and each of three amino acid residues named GLU239, LEU241 and LYS188, were uncovered by molecular docking and molecular dynamics simulation. The cell-based assays showed that compound **Q17** could protect SH-SY5Y cells from okadaic acid (OA)-induced injury by targeting DYRK1A. More importantly, compound **Q17** significantly improved cognitive dysfunction in 3×Tg-AD mice, ameliorated pathological changes, and reduced the expression of DYRK1A, GSK-3β and GSK-3β (pSer9), attenuated tau hyperphosphorylation and Aβ deposition as well. In summary, our computational modeling strategy is effective to identify novel chemotypes of DYRK1A inhibitors with great potential to treat AD, and the identified compound **Q17** in this study is worthy of further study.

**Graphic Abstract:** 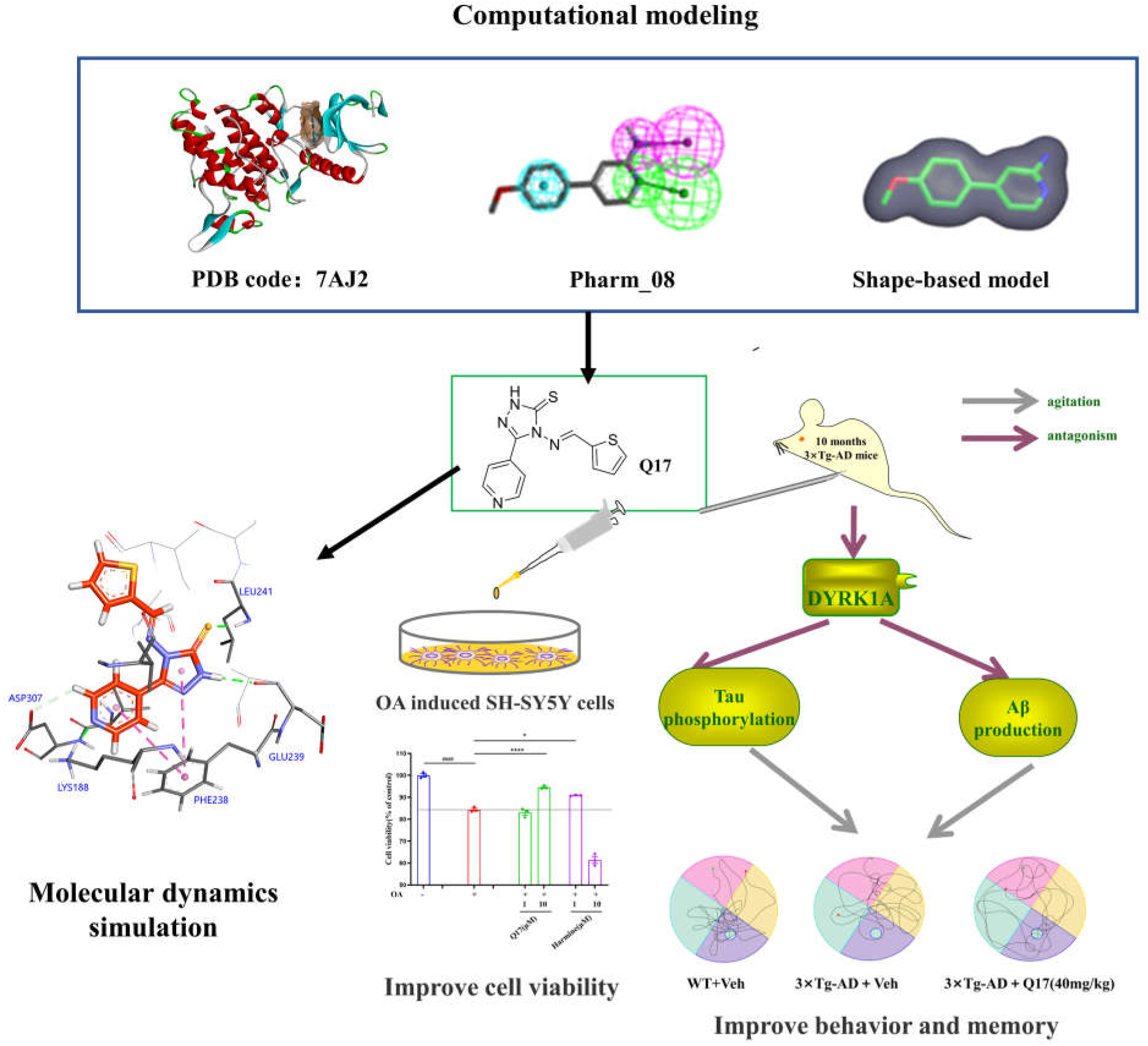

## 1 Introduction

Alzheimer’s disease (AD) is a neurodegenerative disease with age-related cognitive, intellectual, memory and behavioral decline [1, 2]. According to the epidemiology, 50 million people around the world are affected by AD at present [3]. Up to now, marketed drugs for AD only include Donepezil [4] as a central acetylcholinesterase (AChE) inhibitor, and Memantine Hydrochloride [5], an N-methyl-D-aspartic acid (NMDA) receptor antagonist. These drugs could only alleviate symptoms, and the long-term use may lead to severe side effects. Therefore, there is an urgent need to discover and develop new anti-AD drugs.

In the currently known AD pathology, the formation of amyloid β-protein (Aβ) plaques [6] and the hyperphosphorylation of Tau proteins in brain [7] are two important factors. Aβ is mainly derived from the splicing of β-amyloid precursor protein (APP), which is catalyzed by β-secretase and γ-secretase [8]. Phosphorylation of APP leads to the enhanced splicing of APP and the increased Aβ secretion. Tau protein mainly exists in axons, and its normal phosphorylation helps maintain stability of axonal tubulin in brain [9, 10]. But hyper-phosphorylated Tau protein destabilizes tubulin and makes it aggregate into neurotoxic neuronal fiber tangles (NFTs) [11, 12], which eventually results in AD.

The phosphorylation of APP, γ-secretase or Tau protein associated with AD pathology [13] is regulated by dual-specificity tyrosine-(Y)-phosphorylation regulated kinase 1A (DYRK1A) [14]. DYRK1A, belonging to the CMGC family of serine/threonine kinases (i.e. CLKs, MAPKs, GSK-3, CDKs), is one of the conserved DYRK kinases (DYRK1A, DYRK1B, DYRK2, DYRK3 and DYRK4) [15]. DYRK1A is a dual-specificity protein kinase that could phosphorylate itself at tyrosine (Tyr) residues of the activation loop and catalyzes phosphorylation of serine (Ser) and threonine (Thr) residues of its substrates [16]. It has been reported that elevated expression of DYRK1A is observed in different regions of the brain in AD patients [17]. DYRK1A may phosphorylate Thr668 of APP and promote the production of Aβ. It also phosphorylates γ-secretase and enhances its proteolytic activity leading to the increase of Aβ [18]. DYRK1A phosphorylates multiple Sers (S199, S202, S396, S400, S404 and S422) and Thrs (T181, T205, T212, T217 and T231) of Tau protein [19, 20]. Accordingly, the inhibition of DYRK1A may be a potential therapy for AD.

To date, many DYRK1A inhibitors have been identified, but few of them have anti-AD therapeutic effects and none of them has been approved for AD (Fig. 1). Epigallocatechin gallate (EGCG) is currently in the phase II clinical trial. As reported, EGCG improves memory, and is well tolerated by AD patients [21, 22], but it lacks specificity and low bioavailability. SM07883 [7] is a highly active DYRK1A inhibitor. Though its chemical structure has not been disclosed, it is now in phase I clinical trial. Harmine is a well-known DYRK1A inhibitor, with the alkaloid as its core scaffold [23]. Unfortunately, it has not entered clinical trials due to its hallucinogenic effects [24–26]. To the best of our knowledge, the lack of diverse chemotypes somewhat hinders the development of DYRK1A inhibitors as anti-AD drugs.

In this study, we aim to discover a novel chemotype of DYRK1A inhibitors for AD treatment via computational modeling and VS, *in vitro* and *in vivo* biological evaluation. By retrospectively analyzing ligand enrichment of structure-based VS strategies with molecular docking against 70 DYRK1A crystal structures, we identified the well-performing strategy that was based on the crystal structure with the PDB entry of 7AJ2. To speed up VS, we used the selected co-crystal structure (PDB code: 7AJ2) to generate a pharmacophore model and a shape model for ligand-based VS. Subsequently, with the combined strategy of ligand-based and structure-based VS, we were able to select and purchase 16 commercially available potential hits. Compound **Q17** was identified as a novel DYRK1A inhibitor by enzymatic assay and its binding mode to DYRK1A was proposed by molecular dynamics (MD) simulation. The cell-based assay confirmed its neuroprotective activity for the okadaic acid (OA)-induced SH-SY5Y cells. The behavioral study as well as the assays including Hematoxylin–eosin (HE) Staining, western blotting, and immunohistochemistry analysis further demonstrated compound **Q17** could alleviate cognitive dysfunction of 3×Tg-AD mice via reducing the expression of DYRK1A, GSK-3β and attenuating Tau protein phosphorylation and Aβ deposition.

## 2 Materials and Methods

### 2.1 Computational modeling

#### 2.1.1 Protein structure selection: screening power of molecular docking

The method of ligand enrichment (or screening power) assessment was the same as our previous work [31, 32], except for the use of our recently developed tool, MUBD-DecoyMaker 2.0 [33] (https://github.com/jwxia2014/MUBD-DecoyMaker2.0) to generate a benchmarking dataset for DYRK1A (MUBD-DYRK1A). Briefly, the DYRK1A inhibitors from ChEMBL29 and the ZINC library were used as the input of MUBD-DecoyMaker 2.0. Physicochemical property matching, ROC curves from similarity search based on six physicochemical properties or topology (MACCS fingerprints) [34] were used to evaluate the quality of the benchmarking dataset.

Next, 70 available co-crystal structures of *homo sapiens* DYRK1A were retrieved from PDB (https://www.rcsb.org/), cleaned by Discovery Studio (DS, v16.1.0, Dassault Systèmes Biovia Corp.), and superimposed by the module of “Superimpose Proteins” of DS to check the consistency of the binding sites.

Flipper (version 3.1.1.2, OpenEye Scientific Software, Santa Fe, NM) and OMEGA (version 3.1.1.2, OpenEye Scientific Software, Santa Fe, NM) [35] were used to generate isomers and poses of the MUBD compounds, respectively. FRED (version 3.3.1.2, OpenEye Scientific Software, Santa Fe, NM) was used to carry out molecular docking against each protein structure[36]. The compounds were ranked by the Chemgauss4 score. The scores were used for the generation of the ROC curve. ROC AUC and ROC enrichment at 0.5% of the recovered decoys (ROCE0.5%) were calculated based on the ROC curve [37]. ROCE0.5% values were used to represent ligand enrichments of molecular docking approaches, based on which the optimal approach was determined.

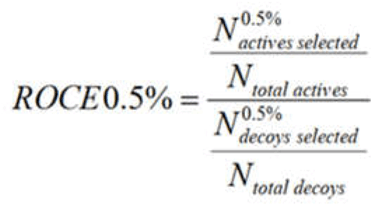

#### 2.1.2 Virtual screening

VS was carried out to identify novel DYRK1A inhibitors from the Specs chemical library (https://www.specs.net/, > 210,000 compounds). To speed up the process of VS, pharmacophore model-based filtering and shape filtering were adopted ahead of molecular docking.

The optimal cocrystal structure of DYRK1A was used for pharmacophore modeling and shape generation. The pharmacophore models were generated, with the "Receptor-Ligand Pharmacophore Generation" module of DS. DYRK1A inhibitors and non-inhibitors were then collected from literature [38] and prepared by the “Prepare Ligand” module. With these compounds to comprise a test set, the "Ligand Profiler" module was applied to validate model performance. Heatmaps were displayed and used for model selection. ROCS (version 3.3.1.2, OpenEye Scientific Software Inc., Santa Fe, NM, USA) was used to generate the shape model based on the binding pose of the cognate ligand in the co-crystal structure [39, 40].

A multi-conformer database of the Specs compounds was built by the “Build 3D Database” module of DS. All the conformers in the database were mapped to the selected pharmacophore model by a rigid fit algorithm (“FAST” search) implemented in the “Search 3D Database” module. The FitValue score was the metric to measure the similarity of each conformer to the pharmacophore. Top-scoring compounds (40%) were kept from this pharmacophore filtering. The shape model was further used for filtering. Shape similarity was measured by the ShapeTanimoto score. Top-scoring compounds (20%) were obtained and submitted for molecular docking with FRED against the optimal receptor structure. Firstly, a maximum of 200 poses for each compound were generated by OMEGA, positioned in the binding site of the receptor and scored by the Chemgauss4 scoring function. 500 top-scoring compounds were kept and their binding modes were visually inspected. The compounds that formed hydrogen bonds with LEU241 and LYS188 were retained. The structural clustering based on the FCFP_6 fingerprints was carried out with the "Cluster Ligands" protocol of DS. By visual inspection, one or two compounds from each cluster were selected by giving priority to those with greater FitValues, ShapeTanimoto scores and Chemgauss4 scores as well as good synthetic feasibility.

#### 2.1.3 Molecular dynamics simulation

Molecular dynamics simulation was performed according to our previously published protocol [32, 41]. The DYRK1A protein topology file was constructed with the GROMOS96 43A1 force field [42], and the ligand topology file was constructed by submitting the PDB coordinates to the LigParGen server (http://zarbi.chem.yale.edu/ligpargen/) [43]. Molecular geometry was optimized based on the theory of HF/6-31G* by Gaussian 09W, the atomic charge was assigned according to the molecular electrostatic potential with the help of the RESP module in the AMBER21 package. Then, GROMACS software (version 2019.4) was used to perform molecular dynamics (MD) simulations of the DYRK1A-ligand complex [44]. The system consisted of 30613 single point charge (SPC) water to solvate the entire system [45]. The water box was then extended by 10 Å from the periphery of the system in each dimension. Afterward, 14 Cl-were added to the system to make the total charge become zero. MD simulation included energy minimization, equilibration, and production phases. The simulation started with 5000 steps of energy minimization based on steepest descent algorithm. In the equilibrium phase, 500 ps simulation for NVT and 500 ps simulation for NPT were included. The system was maintained at a pressure of 1 atm using Parrinello Rahman and a constant temperature of 300 K using V-rescale. Lastly, 200-ns MD simulation without restraint was performed at NPT. The coordinates of the system were saved every 500 ps during the simulation.

### 2.2 Chemical Synthesis

#### 2.2.1 General Methods

All the commercial chemicals and solvents were used without further purification. The ^1^H NMR (500 MHz) spectra were recorded by WNMR-I 500, with tetramethylsilane (TMS) as the internal standard. ^13^C NMR spectra (126 MHz) were recorded by Bruker AVANCEIII 400, with TMS as an internal standard. Low-Resolution Mass Spectrometry was performed by Water ACQUITY QDa mass spectrometer (C18 1.7μm; 2.1×50mm Column).

#### 2.2.2 Procedure for the synthesis of **Q17**

The synthesis was carried out according to the published method [46], with minor modifications. To a 10 mL solution of 1.0 g (7.3 mmol) of isonicotinic acid hydrazide in ethanol was added 0.6 g (11.0 mmol) of potassium hydroxide and 0.6 mL (11.0 mmol) of carbon disulfide. The solution was stirred at room temperature for 30 min. The precipitate produced by filtration was washed with ethanol and dried. The precipitate was then added to 10 mL of water, and then 0.7 mL (14.6 mmol) of hydrazine monohydrate was added. The reaction mixture was heated for 4 h under reflux conditions. After cooling down, the solution was diluted with 10 mL of cold water and acidified with concentrated solution of hydrogen chloride, filtered and driedto afford the intermediate(4-Amino-5-(pyridin-4-yl)-4H-1,2,4-triazole-3-thiol). Using acetic acid as a solvent, 400 mg of the intermediate (2.08 mmol) was refluxed with 400 μL of thiophene-2-carbaldehyde (2.08 mmol) at 110 °C for 8 h. The mixture was left overnight, and then filtered, washed with petroleum ether, and dried to afford the end product.

### 2.3 *In vitro* evaluation

#### 2.3.1 DYRK1A inhibition assay

The assay was carried out in 384-well plates using the ADP-Glo Kinase Assay kit (Promega), according to the published method [47]. To be specific, it was a luminescent ADP detection assay to measure kinase activity by quantifying the amount of ADP produced during a kinase reaction. The kinase reaction between the kinase and its substrate along with the necessary cofactors occurred in the assay buffer composed of Hepes (Thermo), mM MgCl2 (Sigma), Brij35 (Millipore), mM EGTA (Sigma), mM DTT (MCE). Briefly, in the compound, kinase and metal solution, ATP and the substrate solution was added to initiate the reaction. Then, ADP-Glo kinase reagent was added to terminate the reaction, followed by the addition of kinase detection reagent. All of the aforementioned mixtures were incubated at 25°C. The luminescence signal was recorded on a microtiter plate reader. Based on the luminescence, the activity (%) of the DYRK1A protein at different concentrations were determined. GraphPad Prism (GraphPad Software Inc., La Jolla, CA) was used to calculate IC_50_ values. Harmine was used as a positive control. The assay was carried out in triplicate.

#### 2.3.2 siRNA-based DYRK1A knockdown in SH-SY5Y cells

SH-SY5Y cells were cultured with high glucose DMEM medium (KeyGEN, China) containing 10% fetal bovine serum (Hangzhou Sijiqing, China), 1% penicillin and streptomycin, in an incubator at 37 ℃ and with 5% CO_2_. All the cells were sub-cultured for 3 to 5 generations for experiments.

The SH-SY5Y cells were cultured in a 6-well plate, and the transfection experiments were performed when 30%-40% of the cells were fused. siRNA (Fenghbio, China) was diluted with 250 µL of serum-free DMEM medium to a final concentration of 50 nM, and mixed for 5 min. Meanwhile, to another 250 µL serum-free DMEM medium was added 6 µL Lipo8000 (Beyotime, China) and they were mixed for 5 min as well. The two mixtures were kept at room temperature for 30 min to allow full binding of siRNA or Lipo8000, respectively. After 4 h, the serum-contained culture medium was used to ensure cell growth. After transfection for 48 h, siDYRK1A-SH-SY5Y cell model was established.

#### 2.3.3 CCK-8 assay for cell viability

The protocol to evaluate neuroprotective effect on OA-induced SH-SY5Y cells is described as follows. SH-SY5Y cells were seeded in 96-well plates at a density of 5 × 10^3^ per well for 24 h. Then, the cells were treated with or without compounds (dissolved in DMSO, harmine as a positive drug) at the indicated concentrations, and after 24h, 100 nM OA was added to each well. Cell viability was analyzed after 24 h with CCK-8 (Solarbio, China). 10 μL of CCK-8 solution was added to the cells, and then incubated at 37°C for 2 h. The optical density (OD) was measured using a Microplate Reader (TECAN, Switzerland) at 450 nm, and cell viability was expressed as percentage% (compared to negative control). Each assay was repeated at least three times.

#### 2.3.4 Western blot assay

The cells were collected in RIPA lysate (Solarbio, China) containing 1% benzyl sulfonyl fluoride (Solarbio, China) and 1% protein phosphatase inhibitor (Solarbio, China), and then placed on ice for 30 min and further subjected to centrifugation at 4 °C and 12,000 rpm for 15 min. BCA assay (Beyotime, China) was performed for measuring the concentration of protein. Protein samples of equal amount were separated and transferred to the 8-12% SDS-PAGE and PVDF membranes. After being blocked for 2 h at room temperature with 5% non-fat dry milk in TBS-Tween-20 (0.1%, v/v), the membranes were cultured with primary antibody at 4°C overnight, i.e. anti-DYRK1A (1:1000, Abcam, UK), anti-Glycogen synthase kinase 3β (GSK-3β) (1:1000, Abcam, UK), anti-GSK-3β (pSer9) (1:1000, Abcam, UK), anti-Tau (pSer396) (1:1000, Abcam, UK), anti-Tau(1:500, Thermo Fisher Scientific, USA) or anti-β-tublin antibody (1:1000, Zsbio, China). β-tublin was used as the internal control. Subsequently, the membranes were cultured with secondary antibody (1:3000, Abcam, UK) for 1 h at room temperature. The Amersham Imager 600 (GE, USA) was used to measure the blots. Image-J software was used to examine the strip’s grayscale data after film scanning.

### 2.4 *In vivo* evjaluation

#### 2.4.1 Animals and treatment

Triple transgenic (3×Tg-AD) mice [B6; 129-Psen1^tm1Mpm^Tg (APPSwe, tauP301L) 1Lfa/Mmjax] and wild type (WT) mice on mixed C57BL/6JB6 and 129S1/SvImJ background were purchased from The Jackson Laboratory, bred and housed in individually ventilated cages (IVC) of the SPF animal facilities at the Institute of Pharmacology of Shandong First Medical University (Tai’an, China) with a 12-h light/12-h dark cycle (7:00 am-7:00 pm) and food and water ad libitum. 3-4 mice were kept in the same cage and moved to a clean cage every three days. 10-month-old male 3×Tg-AD mice (n=45) were randomly divided into five groups (9 mice per group) and respectively treated by daily intraperitoneal injection of the test compound **Q17** (10mg/kg, 20mg/kg, and 30mg/kg), Harmine (20 mg/kg), vehicle (20% β-cyclodextrin solution) (the “3×Tg-AD” group) for 21 days. WT mice (n=9) were also treated with the 20% β-cyclodextrin solution (the “WT” group). During the period of treatment, the body weight of each mouse was recorded. From day 13 to 21 of administration, all the mice were subjected to various behavioral tests in the following order: the novel object recognition (NOR), Y-maze,

Morris water maze (MWM), and passive avoidance test (PAT). 1 h after the last behavioral test (i.e., the PAT), the mice were killed by decapitation and the brains were collected. The hemispheres of 3 mice in each group were collected for pathological staining. The cerebral cortex and hippocampus of the remaining mice were isolated from the hemispheres for western blot; tissues were stored at −80℃ before use. All the experiments were performed in accordance to the protocols approved by the Institutional Animal Care and Use of Committee of Shandong First Medical University (The experimental animal license was SYXK (Lu) 2017-0023).

#### 2.4.2 NOR test

The task of NOR was performed according to the protocol previously described [48]. The device was an open box with length, width and height as 60cm. The experiment was completed in 3 days: one day for habituation, one day for training, and one day for testing. During training, the mice were allowed to explore 2 identical objects for 5 min. On the day for testing, one of the training objects was replaced with a new object, and the mice were allowed to freely explore the field in the presence of the familiar object and the novel object for 5 min. The Top Scan system (CSI, USA) was used to record and analyze the time the mice spent on sniffing new and familiar objects, and compare each group’s recognition object index. Recognition object index was the ratio of the time spent exploring the novel object to the sum of the time spent on exploring the novel object and the time spent on exploring the familiar object.

#### 2.4.3 Y-maze test

The Y-maze consisted of three arms, each with an angle of 120° and a length of 40 cm. The starting point for the mice was the center of the Y-maze, and the Top Scan system (CSI, USA) was used to record the times for which the mouse went to each arm and calculate the maximum of arm entries and spontaneous alternation [49]. Spontaneous alternation equals to the ratio of alternations to maximum number of arm entries.

#### 2.4.4 MWM test

The MWM was used to test the behavior of mice [50]. The pool was divided into four quadrants, and the platform for escape was located at the fourth quadrant. The water was about 35 cm deep, and the platform was 1.5 cm below water. The water was dyed white by the whitening agent, and the water temperature was maintained at 22 ± 2 °C. On Day 1 to Day 5, each mouse was trained according to the standard protocol consisting of 4 consecutive trials per day. If the platform was found in 60 s, the escape latency of the mice was recorded. If not, the mouse was gently guided to the platform and allowed to stay there for 10 s. The space exploration experiment was arranged on Day 6. Before the experiment, the platform for escape was removed from the maze, and the mice were allowed to swim freely in water for 60 s. The swimming speed, the time to cross the platform, the latency period to enter the platform, and the time to stay in the quadrant of the platform were recorded.

#### 2.4.5 PAT test

The PAT was carried out according to the previous study [51]. The device included two compartments, a bright compartment and a dark compartment in the same shape and size, and an automatic door connecting two compartments. The bottom of the dark compartment was equipped with stainless steel rods connected to an electroshock generator. The protocol included three steps: adaptation, acquisition, and testing. In the adaptation and acquisition, the mice were placed in the bright compartment for 2 min, then the automatic door was opened to allow the mice to enter the dark compartment. As soon as the mice entered the dark compartment, the door was closed and a mild electric shock (0.5mA, 3s, repeated three times) was delivered. After 24 h, the testing was carried out. The automatic door between the light and dark compartments was opened and the dark compartment was not electrified. The mice were placed in the light compartment and allowed to move freely for 2 min. The latency time of the mice to enter the dark compartment was recorded.

#### 2.4.6 HE Staining

The brain tissue was fixed with 4% paraformaldehyde, dehydrated, made transparently, and embedded in paraffin before being sectioned. The paraffin sections were baked for 30 min at 64 ℃ to prevent dewaxing. After dewaxing and rehydration with water, the sections were stained with hematoxylin for 4 min, differentiated with 1% alcohol hydrochloride, and then rinsed with water. The sections were counterstained with eosin for 2 min and dehydrated in ascending grades of ethanol and made transparent with xylene. After that, a light microscope was used to examine the morphological change in the section.

#### 2.4.7 Immunohistochemistry assay

The slices were dried at 60 °C in the oven and stored at room temperature. The slices were deparaffinized, rehydrated, subjected to antigen retrieval, and rinsed in distilled water. The slices were blocked with 5% goat serum (Beyotime, China) for 30 min and then incubated with anti-Aβ_1-42_ (1:300, Abcam, UK), anti-Tau(pSer396) (1:4000, Abcam, UK) and anti-Tau (1:4000, Abcam, UK) antibodies at 4°C overnight. On the next day, the appropriate species secondary antibodies (HRP-conjugated) were added to cover the tissues and incubated for 50 min at room temperature. The slices were then placed in PBS (PH7.4) for decolorization. After the slices were dried, freshly prepared DAB chromogenic solution was dropped into the circle. The color development was checked with the microscope till the color became brown. Then, the double-distilled water was added to stop the color development. Hematoxylin was counterstained for about 5 min, washed with double-distilled water, differentiated with hematoxylin differentiation solution for a few seconds, rinsed again with double-distilled water. When the hematoxylin solution became blue, it was rinsed with the running water. Lastly, the slices were dehydrated and transparent. The slices were removed from toluene and dried, before the slices were covered with neutral gum.

#### 2.4.8 Western blot assay

The western blot assay here was performed in the same way as that in *in vitro* evaluation. Briefly, the tissues were collected, from which the total protein was extracted and the expression of individual protein indicators was determined. Here, the primary antibodies included: anti-DYRK1A (1:1000, Abcam, UK), anti-GSK-3β (1:1000, Abcam, UK), anti-GSK-3β (pSer9) (1:1000, Abcam, UK), anti-APP (1:1000, Cell signaling technology, USA), anti-Presenilin 1 (PS1) (1:5000, Abcam, UK), anti-Aβ_1-42_ (Proteintech, 1:500, China), anti-Neprilysin (NEP) (1:1000, Abcam, UK), anti-Insulin Degrading Enzyme (IDE) (1:1000, Abcam, UK), anti-Tau (pSer396) (1:1000, Abcam, UK), anti-Tau (pSer404) (1:1000, Abcam, UK), anti-Tau(pSer199) (1:1000, Abcam, UK), anti-Tau(pSer202) (1:1000, Abcam, UK), anti-Tau(1:500, Thermo Fisher Scientific, USA) and anti-β-tublin antibody (1:1000, Zsbio, China). β-tublin was used as the internal loading control.

### 2.5 Statistical analyses

Prism 8.4.0 (GraphPad Software Inc, La Jolla, CA, USA) was used for statistical analyses. All the data were expressed as means ± S.E.M of at least three independent experiments. The difference between two normal data was analyzed by using two-tailed t-test, and the between-group statistics was based on ANOVA (SNK-q test was used for pair-by-pair comparisons between groups).

## 3 Results and Discussions

### 3.1 Computational modeling and DYRK1A inhibition assay identified hit compound Q17

#### 3.1.1 The optimal protein structure determined for structure-based VS

By manual curation and visual inspection, 70 co-crystal structures with the cognate ligands located at the same binding site were obtained. To select the protein structure optimal for prospective FRED docking-based VS, we performed retrospective small-scale VS of a benchmarking dataset by FRED docking against different DYRK1A crystal structures and estimated ligand enrichments [27].

MUBD-DYRK1A contains 128 ligands and 4992 decoys (https://github.com/jwxia2014/DYRK1A-Dataset). Fig. 2A shows that six physicochemical properties of ligands and decoys matched well, while Fig. 2B indicates that the ligands and decoys were difficult to differentiate merely by similarity search based on six physicochemical properties or topology (MACCS fingerprints). According to Fig. 2C, molecular docking with FRED against the protein structure with the PDB code of 7AJ2 showed the highest enrichment in term of ROCE0.5%. The exact values of ROCE0.5% for all the SBVS approaches are listed in Table S1, and the value for 7AJ2 is 50.0. Fig. 2D shows the details of protein-ligand interactions from the selected cocrystal structure (PDB code: 7AJ2). The cognate ligand interacts with LYS188 and LEU241 in the form of hydrogen bond, and its benzene ring and pyridine ring form hydrophobic interactions with Val173, Ala186, Leu294 and Val306.

**Fig.1.**
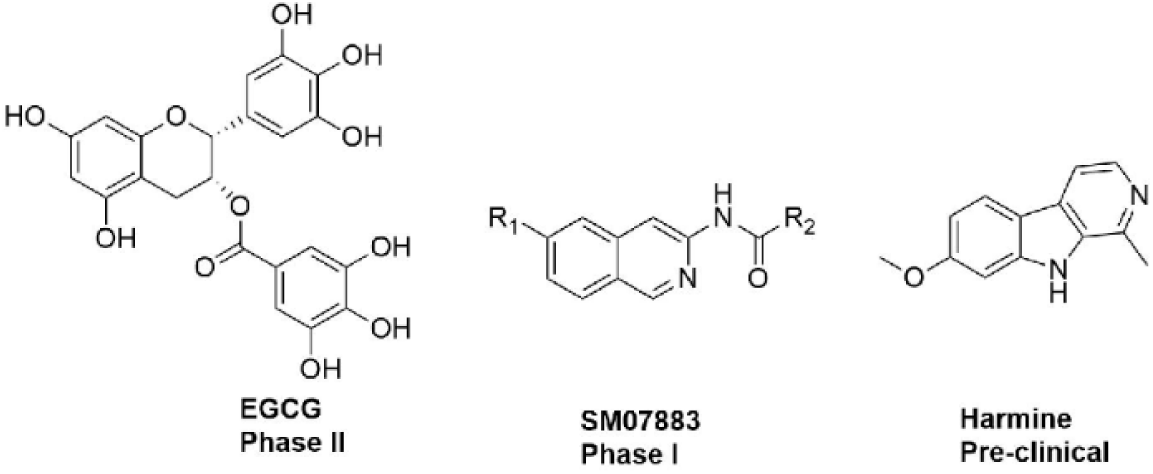
Representative DYRK1A inhibitors that have been tested for Alzheimer’s disease Virtual screening (VS) allows enrichment of the most promising compounds for bioassay in a relatively short period of time [27]. Several research groups have identified DYRK1A inhibitors from large-scale chemical library by using VS. Early in 2012, Wang *et al*. performed homology model-based VS and discovered two micromolar-level DYRK1A inhibitors that were more potent than EGCG [28]. Koyamaet *et al*. constructed a logistic regression model based on residue binding free energy and a pharmacophore model to virtually screen for DYRK1A inhibitors and identified two selective DYRK1A inhibitors with IC_50_ values at the micromolar level [29]. More recently, Choi *et al*. identified aristolactam BIII as a novel DYRK1A inhibitor that could rescue Down syndrome-related phenotypes *in vitro* and *in vivo* via structure-based VS [30]. The aforementioned case of success demonstrates the power of *in silico* methods in discovery of DYRK1A inhibitors, and also indicates different VS methods may contribute to discovery of diverse scaffolds.

**Fig. 2.**
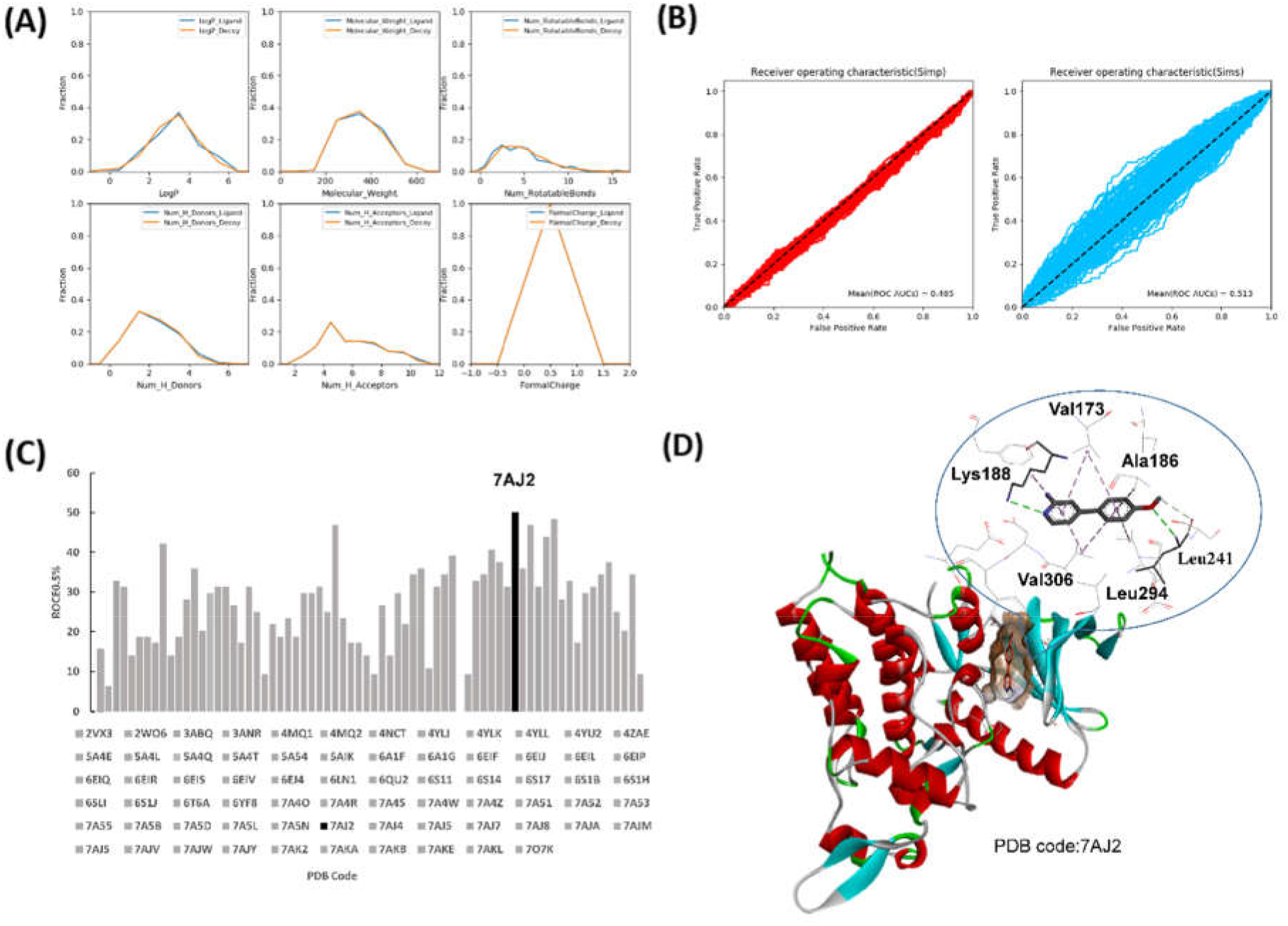
Ligand enrichment assessment of virtual screening approaches using molecular docking with FRED against different DYRK1A protein structures. (A) Physicochemical property matching between MUBD-DYRK1A ligands and decoys. (B) ROC curves from similarity search based on six physicochemical properties or topology (MACCS fingerprints). (C) The ligand enrichment in term of ROCE0.5%. The crystal structure of the highest enrichment (PDB code: 7AJ2) is highlighted. (D) The crystal structure of DYRK1A in complex with its cognate ligand (PDB code: 7AJ2). The binding mode is depicted by Discovery Studio 2016.

#### 3.1.2 The workflow of VS and potential hits

As we mentioned in the Method section, we generated a pharmacophore model and a shape model as the filters ahead of FRED docking in order to speed up VS. The 10 pharmacophore models that were generated by the Catalyst/HipHop module of DS are shown in Fig. S1A. The compounds (test set) used to evaluate the pharmacophore models for their power to differentiate between active (IC_50_<500nM) and inactive compounds (IC_50_>500nM) are shown in Fig. S1B. According to Fig. S1C, it was easy to identify **Pharm_08** as the optimal model (Fig. 3A), as it assigned the maximal values to the active compounds and the minimal values to the inactive compounds. This model was composed of one hydrogen bond acceptor, one hydrogen bond donor and one hydrophobic feature. The shape model generated by ROCS is shown in Fig. 3A.

**Fig. 3.**
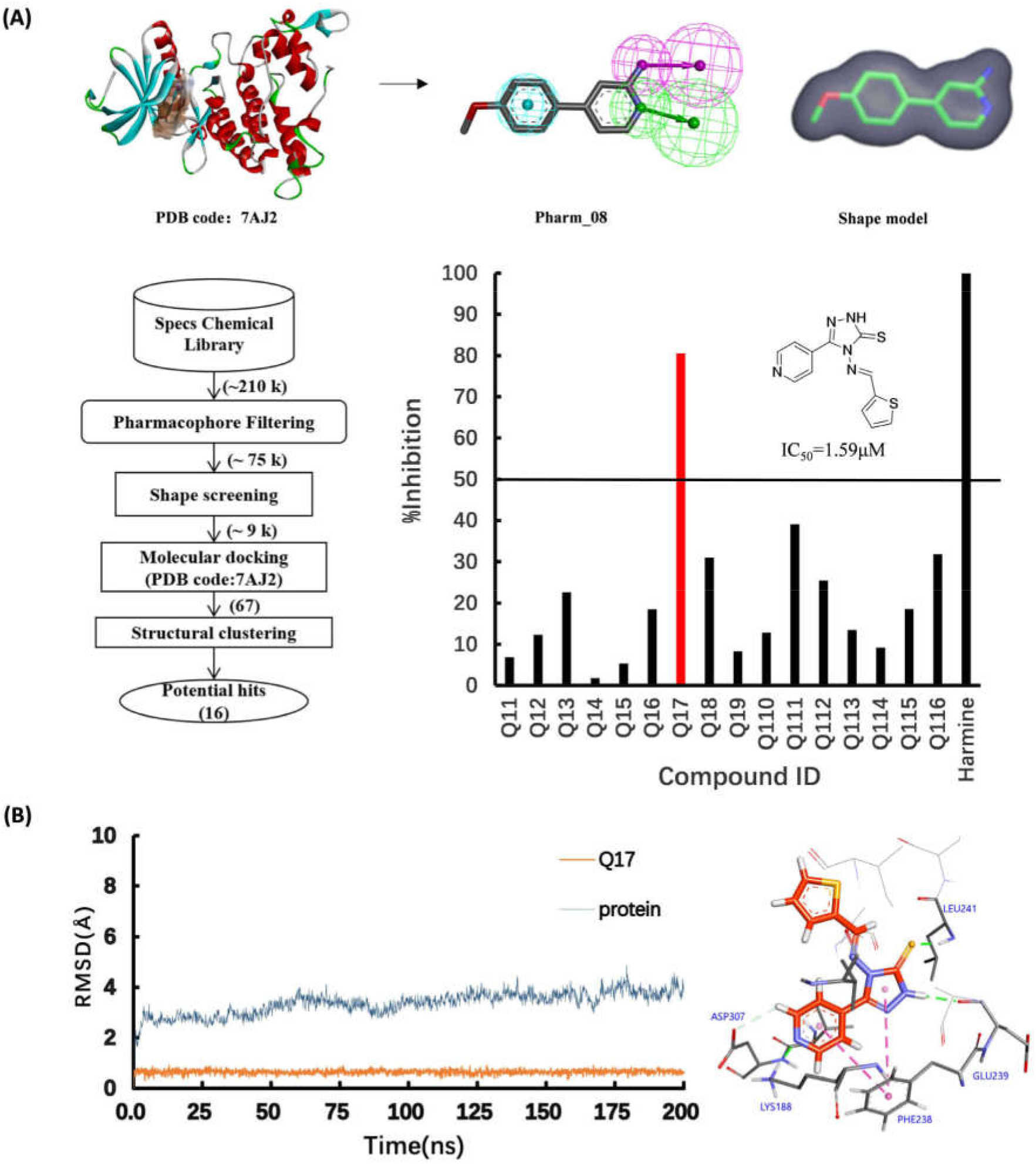
Identification of **Q17** as a novel DYRK1A inhibitor by virtual screening and experimental validation. (A) The crystal structure used for computational modeling (PDB code: 7AJ2), pharmacophore model (Pharm_08) and shape model, the computational workflow and the potential hits (including **Q17**) and their DYRK1A inhibition rate (%) at 10 µM. (B) Molecular dynamic simulation to uncover the binding modes of **Q17** to DYRK1A, including the heavy-atom RMSDs of the DYRK1A protein and **Q17** during 200 ns MD simulation and the detailed binding mode.

We used the VS workflow to screen the Specs chemical library (∼210,000 compounds, Fig. 3A). About 75,000 compounds with the FitValue values greater than 2.5 were retained. 9000 top-scoring compounds in term of ShapeTanimoto score were selected. From molecular docking against the protein structure (PDB code: 7AJ2), top 500 compounds with FRED Chemgauss4 scores less than -11.6075 were selected. According to the binding modes, 67 compounds were retained. 20 clusters were generated by the structural clustering based on FCFP_6 fingerprints. At last, 20 diverse compounds were selected. At that time, only 16 compounds were available to purchase (Q11-Q116, Fig. S2). Their FitValue, ShapeTanimoto and FRED Chemgauss4 score are listed in Table S2.

### 3.1.3 Hit compound Q17: chemical structure, DYRK1A inhibition and binding mode

The DYRK1A inhibition at the concentration of 10 µM is shown in Fig. 3A. Among them, **Q17** (AO-476/40829177) showed the inhibition rate of 80.55%. Its IC_50_ of **Q17** was further determined as 1.59 μM, indicating it was a moderately potent DYRK1A inhibitor. We further performed 200 ns molecular dynamics (MD) simulation to study the binding of **Q17** to the DYRK1A protein. As for the DYRK1A-**Q17** system, it reached equilibrium after about 110 ns (Fig. 3B). The protein-ligand binding conformations after equilibrium were extracted and analyzed. According to Fig. 3B, compound **Q17** formed hydrogen bonds with LEU241 and GLU239 through its core scaffold, a hydrogen bond with LYS188 via the pyridine fragment, and π-π stacking with PHE238.

### 3.2 Compound Q17 exerted neuroprotective activity by targeting DYRK1A

Based on the previous method [52], compound **Q17** was further evaluated for its neuroprotective activity from OA-induced toxicity to SH-SY5Y cells. As shown in Fig. 4A, OA significantly decreased the viability of SH-SY5Y cells at the concentration of 100 nM. Though compound **Q17** was not active at 1 μM, it showed neuroprotective effect at the concentration of 10 μM (Fig. 4A). Interestingly, we observed the positive drug, Harmine at 1 μM but not 10 μM could ameliorate the injury induced by OA. We hypothesized this observation may be attributed to the cytotoxicity of Harmine itself to SH-SY5Y cells at 10 μM. Fig. S3 validated our hypothesis. SH-SY5Y cell viability was significantly decreased by Harmine at the concentration of 10 μM. As for compound **Q17** that was tested in the same assay, it had no effect on SH-SY5Y cells at both 10 μM and 1 μM. This ruled out the bias of compound **Q17** itself on cell viability and thus further confirmed its neuroprotective effect on SH-SY5Y cells.

**Fig. 4.**
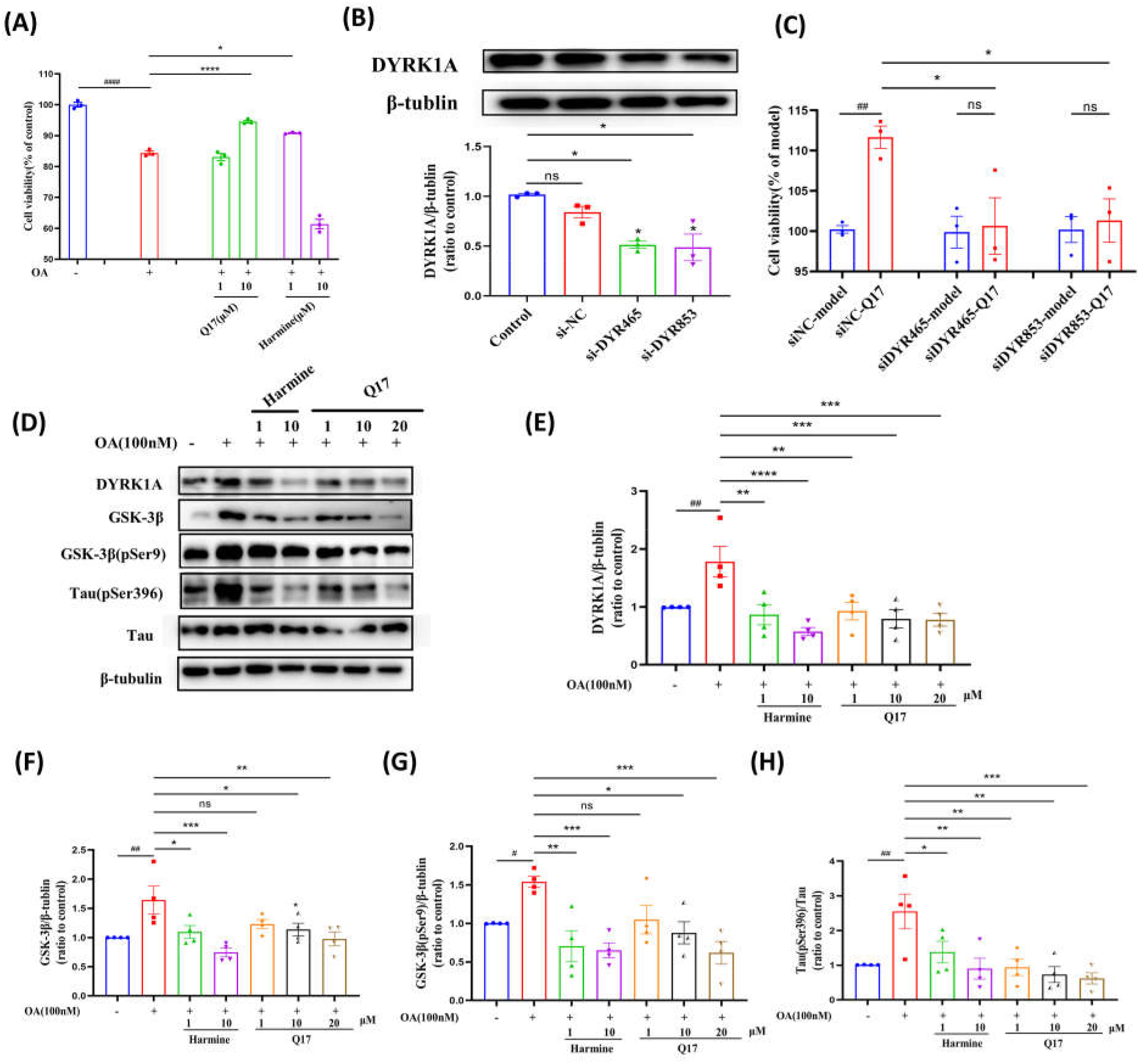
Compound **Q17** exerted neuroprotective activity by targeting DYRK1A. (A) The effect of compound **Q17** on OA-induced SH-SY5Y cells. (B) DYRK1A protein expression in the SH-SY5Y cells treated by siRNA. (C) The effect of compound **Q17** (10 μM) on OA-induced SH-SY5Y cells after siRNA-based knockdown of DYRK1A. The cell viability was set as 100% for the normal cells. The results are presented as mean ± SEM (n = 3). ^##^ *p*< 0.01 vs siNC-model group. * *p* < 0.05 vs siNC-**Q17** group. (D) The effect of compound **Q17** on OA-induced protein expression in SH-SY5Y cells. Representative images are shown. (E-H) Quantification of DYRK1A expression (E), GSK-3β expression (F), GSK-3β (pSer9) expression (G) and Tau (Ser396)/Tau(H). The results are presented as mean ± SEM (n = 4). ^#^*p* < 0.05, ^##^ *p* < 0.01 vs the control group. * *p* < 0.05, ** *p* < 0.01, *** *p* < 0.001, **** *p* < 0.0001 vs the OA (100 nM) group. Harmine was used as a positive drug.

To prove that the neuroprotective effect of compound **Q17** was from the targeting of DYRK1A, siRNA was used to knock down the expression of DYRK1A in SH-SY5Y cells and the effect of compound **Q17** was tested on this cell model. After siRNA-based knock down, the expression of DYRK1A significantly decreased compared with the control group (Fig. 4B), indicating the siDYRK1A-SH-SY5Y cell model was successfully established. Normally, compound **Q17** at 10 μM could improve cell viability of the OA-induced SH-SY5Y cells. Unfortunately, the protective effect of compound **Q17** was significantly weakened when it was tested on the siDYRK1A-SH-SY5Y cell model (Fig. 4C).

It is well known that the onset of AD causes a significant increase in DYRK1A protein expression, which acts synergistically with GSK-3β to induce tau protein phosphorylation. Therefore, it is natural to measure the expression of those proteins in SH-SY5Y cells. The representative bands from the western blot assay are shown in Fig. 4D. As expected, the DYRK1A protein expression level in the OA-induced SH-SY5Y cell model was higher than that in the normal SH-SY5Y cells. Consistently, the levels of GSK-3β protein, pGSK-3β protein and the Tau (pSer396)/Tau also increased, compared with the normal cells. Interestingly, similar to the positive drug, Harmine (1 and 10 μM), compound **Q17** (1, 10, and 20 μM) could reduce the levels of DYRK1A protein, GSK-3β protein, GSK-3β (pSer9) protein and Tau (pSer396)/Tau that had been enhanced in the OA-induced SH-SY5Y cells (Fig. 4F-H).

### 3.3 *In vivo* efficacy for AD

#### 3.3.1 Compound Q17 attenuated cognitive impairment in 3×Tg-AD mice

To evaluate efficacy of compound **Q17** for AD, we carried out chemical synthesis of the compound and obtained the amount sufficient for *in vivo* study (cf. Scheme S4 and methods section for details). We tested compound **Q17** on 3×Tg-AD mice according to the protocol described in Fig. 5A. In general, the *in vivo* experiments included the behavioral study (the NOR test, the Y-maze test, the MWM test and the PAT test), pathological examination and molecule-level assays.

**Fig. 5.**
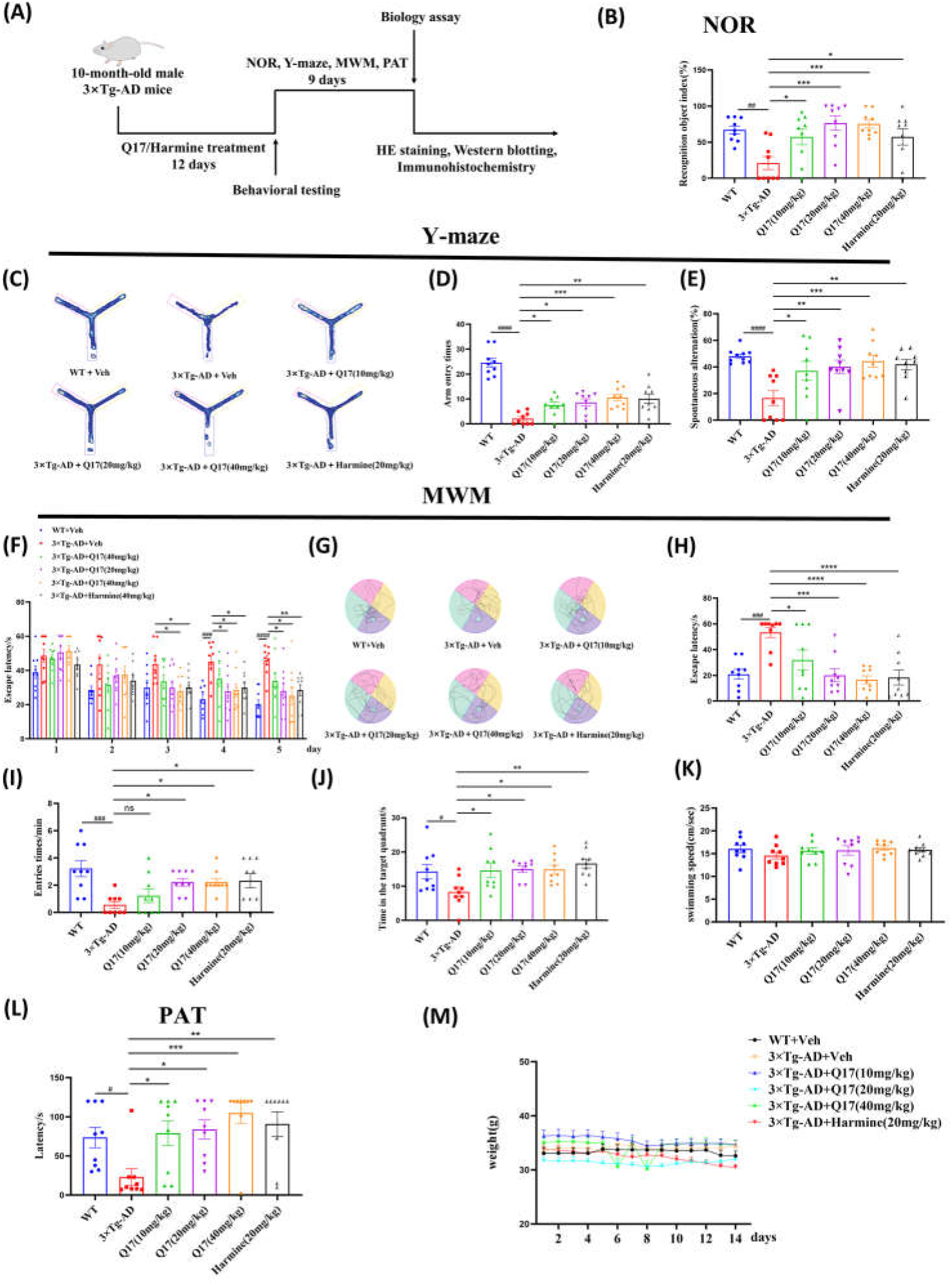
Compound **Q17** improved learning and memory of 3×Tg-AD mice. (A) Protocol of the *in vivo* experiments. (B) NOR test: recognition object index. (C-E) Y-maze test: representative searching paths of the mice (C), arm entry times (D) and spontaneous alternation (E). (F-K) MWM test: the escape latency in the positioning navigation experiments (F), representative swimming paths of the mice (G), the escape latency (H), times of entry to the platform (I), the time spent in the target quadrant (J), the swimming speed of the mice (K). (L) Escape latency in PAT test. (M) Body weight of the mice within 14 days of intraperitoneal administration. The results are presented as mean ± SEM (n = 9). ^#^*p* < 0.05, ^##^ *p* < 0.01, ^###^ *p*< 0.001, ^####^ *p*< 0.0001 vs WT group. * *p* < 0.05, ** *p* < 0.01, *** *p* < 0.001, **** *p* < 0.0001 vs 3×Tg-AD group. Harmine was used as a positive drug.

In the NOR test, the NOR index of the 3×Tg-AD group was less than the WT group (Fig. 5B). In the Y-maze test, the arm entry times and spontaneous alternation of 3×Tg-AD group was significantly lower as well (Fig. 5C-E). The MWM test also showed that the escape latency of 3×Tg-AD mice became longer, as the duration of the positioning navigation experiment and the learning impairment in the space exploration experiment increased (Fig. 5F-J). In the PAT test, the latency of 3×Tg-AD mice decreased as well (Fig. 5L). The positive drug, Harmine, has been reported to have good anti-AD activity in vivo [53]. Consistently, Harmine in this study was observed to improve the aforementioned AD-like behaviors (Fig. 5B-L). The above data indicated that 3×Tg-AD mice was in the status of cognitive impairment thus suitable for evaluating anti-AD efficacy of compounds.

Encouragingly, compound **Q17** could (1) improve the NOR index in NOR test (Fig. 5B), (2) increase the arm entry times and spontaneous alternation in Y-maze test (Fig. 5C-E), (3) reduce the escape latency and weaken the spatial learning impairment, lead to shorter escape latency, more times of entry to the platform and longer time spent in the target quadrant in MWM test (Fig. 5F-J), and (4) boost the latency time in the PAT test (Fig. 5L). We also measured the swimming speed among the six groups, and found there was no significant difference thus excluded the likeliness that athleticism or the other health problems may affect the behavioral study (Fig. 5K). These results demonstrated compound **Q17** was effective to treat 3×Tg-AD mice. In order to preliminarily evaluate the safety profile of the compound on mice, we measured body weight of the mice every day during the 14-day treatment. It was exciting to see that the body weight was not significantly affected by compound **Q17** at all the doses (Fig. 5M).

#### 3.3.2 Compound Q17 attenuated pathological lesions in 3×Tg-AD mice

To explore how compound **Q17** improved behavior in 3×Tg-AD mice, we examined pathological changes in brain regions of the mice by the HE staining assay. By comparing the pathology of 3×Tg-AD mice and WT mice, it was obvious to see that the cytoplasm of the neurons, the demarcation with the surrounding tissues in the hippocampus and cortex became vague. In addition, the neurons were not neatly arranged and the nucleoli were obviously solidified. After the administration of compound **Q17**, the nucleoli of the neurons were visible, and the solidification decreased (Fig. 6). This suggested compound **Q17** could ameliorate pathological lesions in the brain of AD mice.

**Fig. 6.**
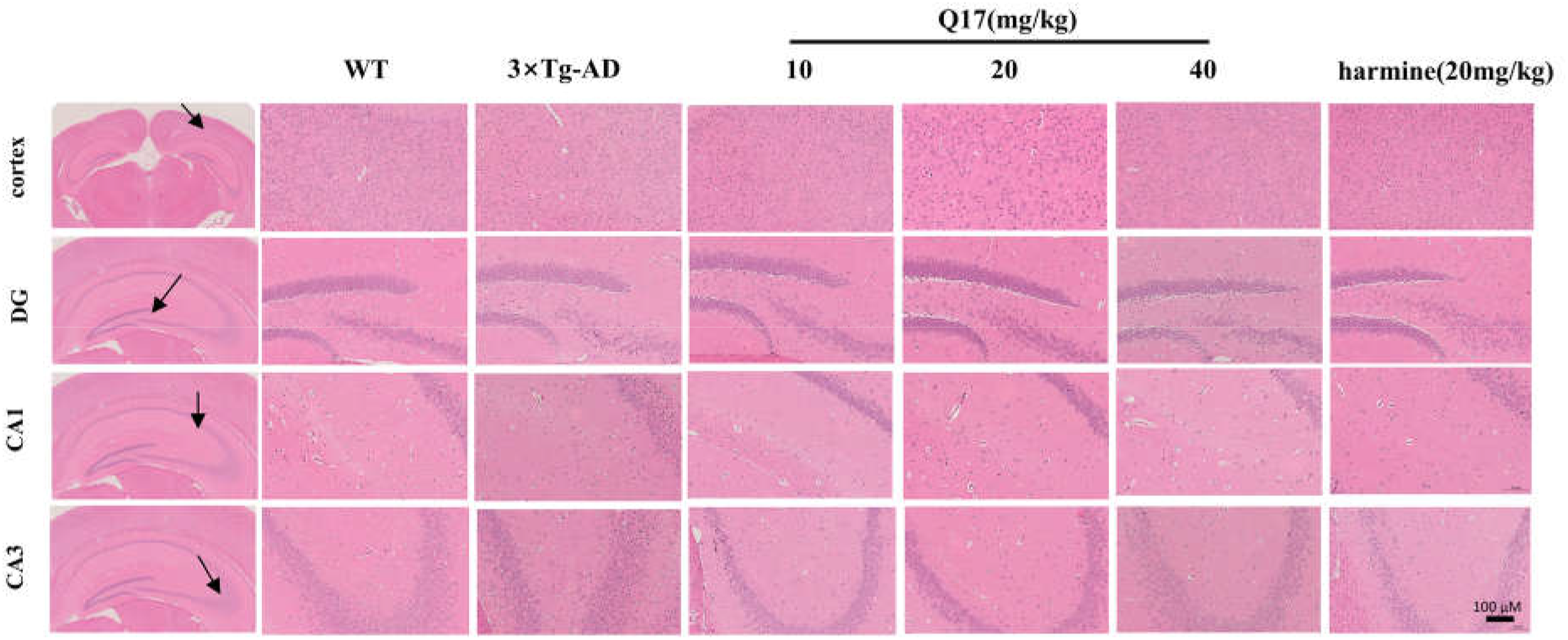
Compound **Q17** attenuated the pathological abnormality in 3×Tg-AD mice. The pathology of 3×Tg-AD mice in the cortex and hippocampus (DG, CA1, CA3). Scale bar represents 100 μm. Harmine was used as the positive drug.

#### 3.3.3 Compound Q17 reduced the phosphorylation of Tau protein in 3×Tg-AD mice

In 3× Tg-AD mice, phosphorylated tau protein was measured by both immunochemistry and western blot assay, as it is acknowledged as a key pathological feature of AD [54]. It was noted that the expression of Tau (pSer396) in the cortex and hippocampus (DG, CA1, CA3 regions), and Tau (pSer396), Tau (pSer404), Tau (pSer199), Tau (pSer202) in the cortex and hippocampus increased in 3×Tg-AD mice, compared with WT mice (Fig. 7A-F, Fig. 8A-B). After the administration of compound **Q17,** the phosphorylated Tau proteins in the cortex and hippocampus were significantly reduced in a dose-dependent manner (Fig. 7A-F, Fig. 8A, B).

**Fig. 7.**
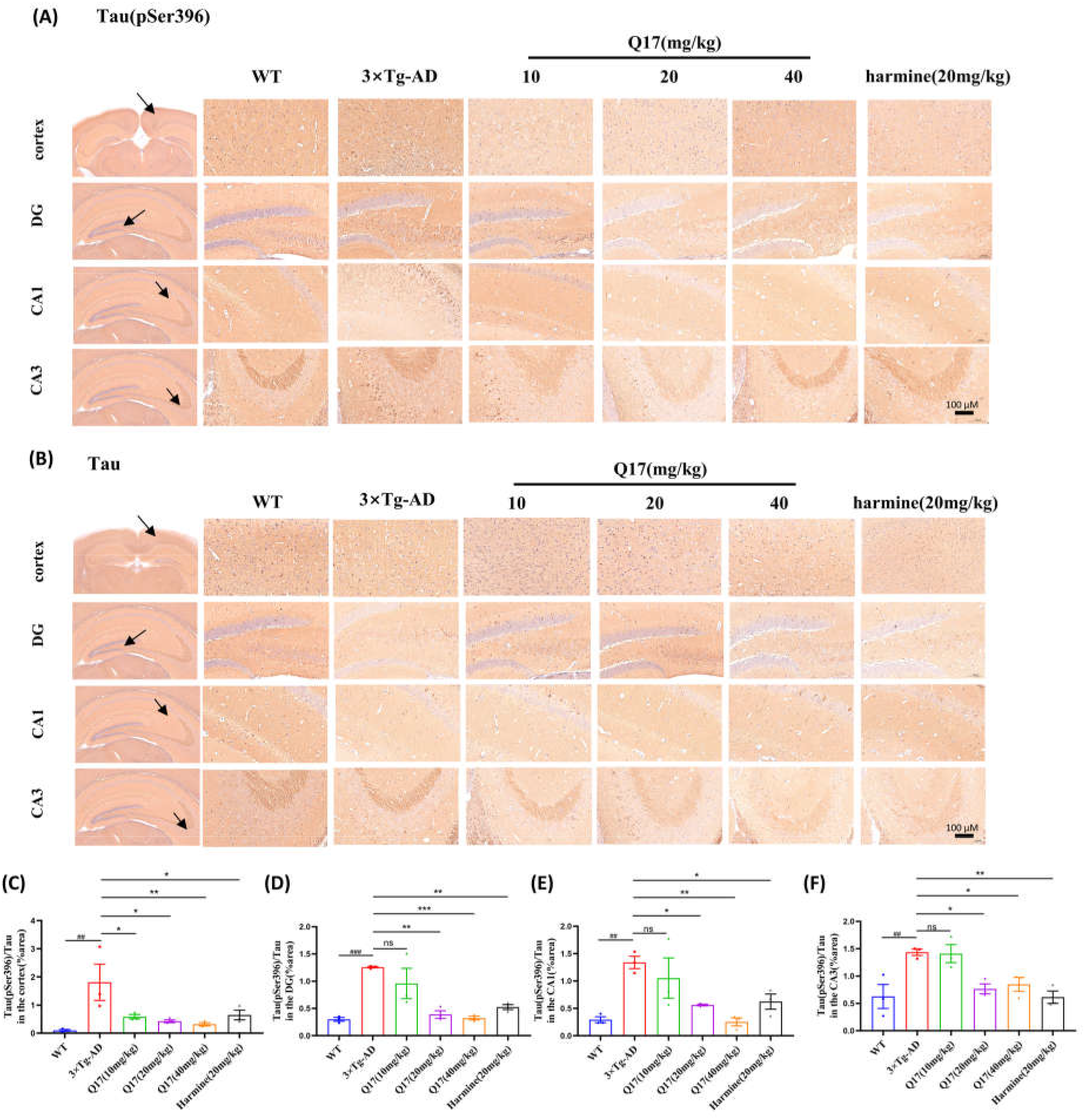
Compound **Q17** reduced tau protein phosphorylation of 3×Tg-AD mice, shown by the immunochemistry assay. (A-B) The effect of compound **Q17** on the tau protein phosphorylation in the cortex and hippocampus (DG, CA1, CA3) of 3×Tg-AD mice. (C-F) Quantification of Tau(pSer396)/Tau by immunochemistry in the cortex (C) and hippocampus (DG, CA1, CA3) (D-F). Scale bar represents 100 μm. Bars shown are means ± SEM (n=3). ^##^*p* < 0.01, ^###^*p* < 0.001 vs WT group. \**p* < 0.05, \*\**p* < 0.01, \*\*\**p* < 0.001 vs 3×Tg-AD group. Harmine was used as the positive drug.

**Fig. 8.**
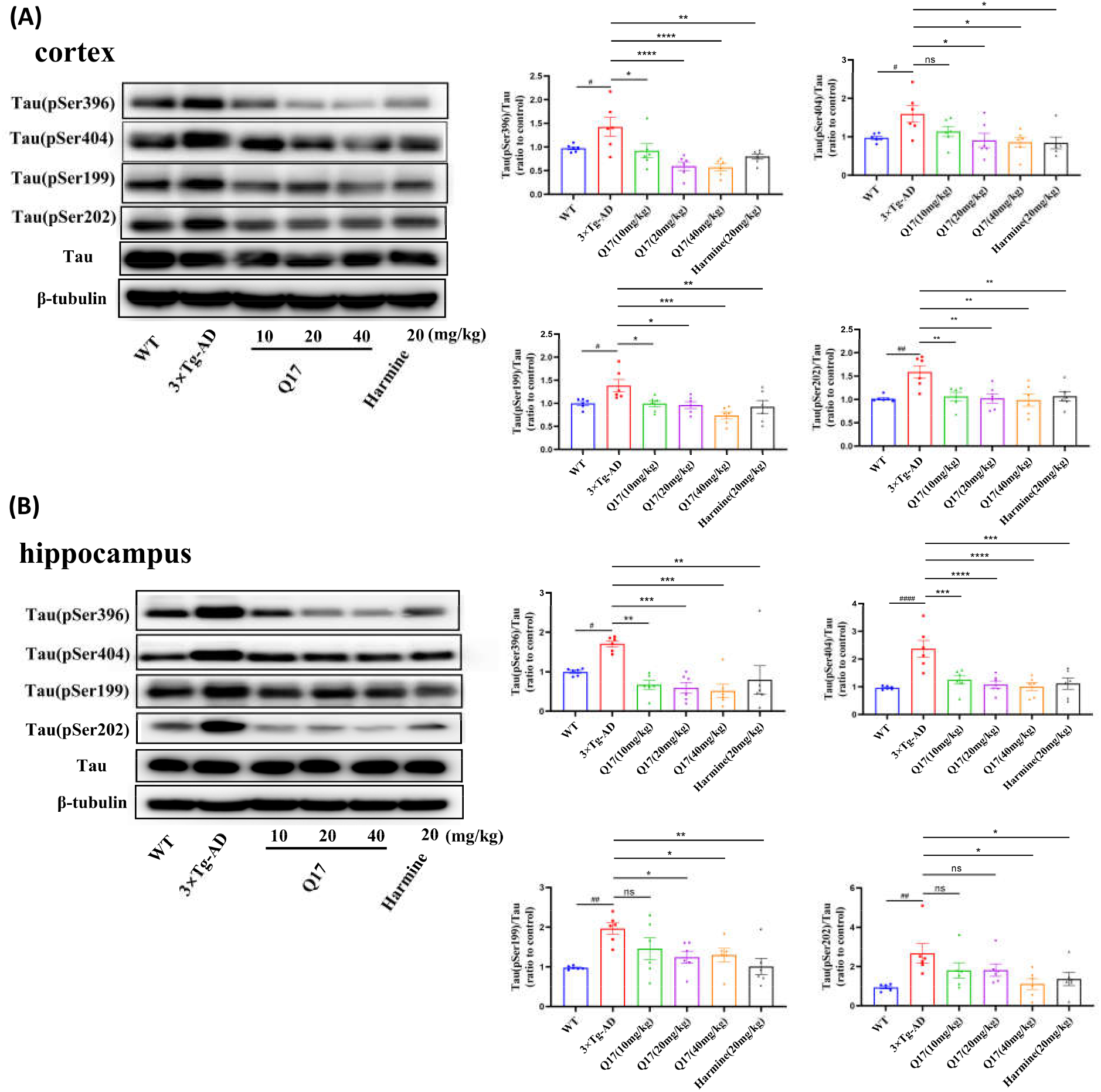
Compounds **Q17** reduced tau protein phosphorylation of 3×Tg-AD mice, shown by the western blot assay. The effect of **Q17** on Tau protein phosphorylation in the cortex (A) and hippocampus (B). Representative images are shown. The levels of Tau (pSer396) expression, Tau (pSer404) expression, Tau (pSer199) expression and Tau (pSer202) expression in the cortex and hippocampus were analyzed. The results are presented as mean ± SEM (n = 6). ^#^*p* < 0.05, ^##^*p* < 0.01, ^####^*p*< 0.0001 vs WT group. \**p* < 0.05, \*\**p* < 0.01, \*\*\**p* < 0.001, \*\*\*\**p* < 0.0001 vs 3×Tg-AD group. Harmine was used as a positive drug.

#### 3.3.4 Compound Q17 delayed the development of Aβ plaque deposition in 3×Tg-AD mice

As one of the key pathological features of AD development, Aβ deposition was also embodied in 3× Tg-AD mice [55, 56]. Accordingly, we measured the Aβ deposition-related indicators by using both immunochemistry and western blot assay. As shown in Fig. 9A-E and Fig. 10A-B, the expression of APP, PS1, Aβ_1-42_ in the cortex and hippocampus (DG, CA1, CA3 regions) of 3×Tg-AD mice, significantly increased, compared with WT mice. After compound treatment, **Q17** (20 mg/kg and 40 mg/kg) reduced Aβ deposition-related indicators in both cortex and hippocampus in a dose-dependent manner. We also observed that **Q17** did not significantly improve the expression of Aβ degradation-related proteins NEP and IDE in the brain, suggesting that **Q17** may not be involved in the Aβ degradation pathway (Fig. 9A-E, Fig. 10A-B). This is advantageous because toxicity caused by over-Aβ degradation in central nervous system may not exist for compound **Q17** [57].

**Fig. 9.**
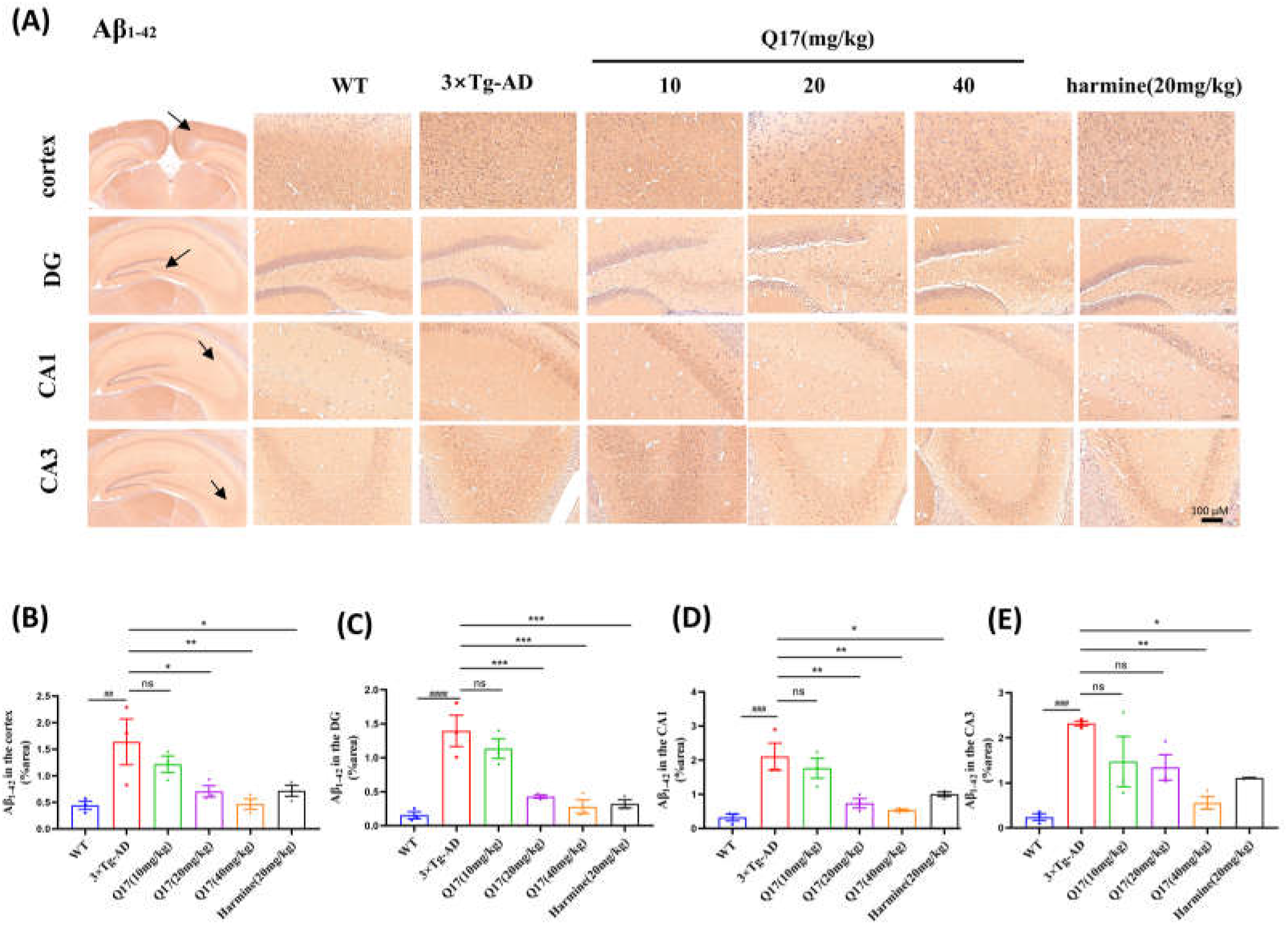
Compound **Q17** reduced Aβ deposition of 3×Tg-AD mice, shown by the immunochemistry assay. (A) The effect of compound **Q17** on the Aβ deposition in the cortex and hippocampus (DG, CA1, CA3) of 3×Tg-AD mice. (B-E) Quantification of Aβ_1–42_ by H&E staining in the cortex (B) and hippocampus (DG, CA1, CA3) (C-E). Scale bar represents 100 μm. Bars shown are means ± SEM(n=3). ^##^*p* < 0.01, ^###^*p* < 0.001, ^####^*p*< 0.0001 vs WT group. \**p* < 0.05, \*\**p* < 0.01, \*\*\**p* < 0.001 vs 3×Tg-AD group. Harmine was used as a positive drug.

**Fig. 10.**
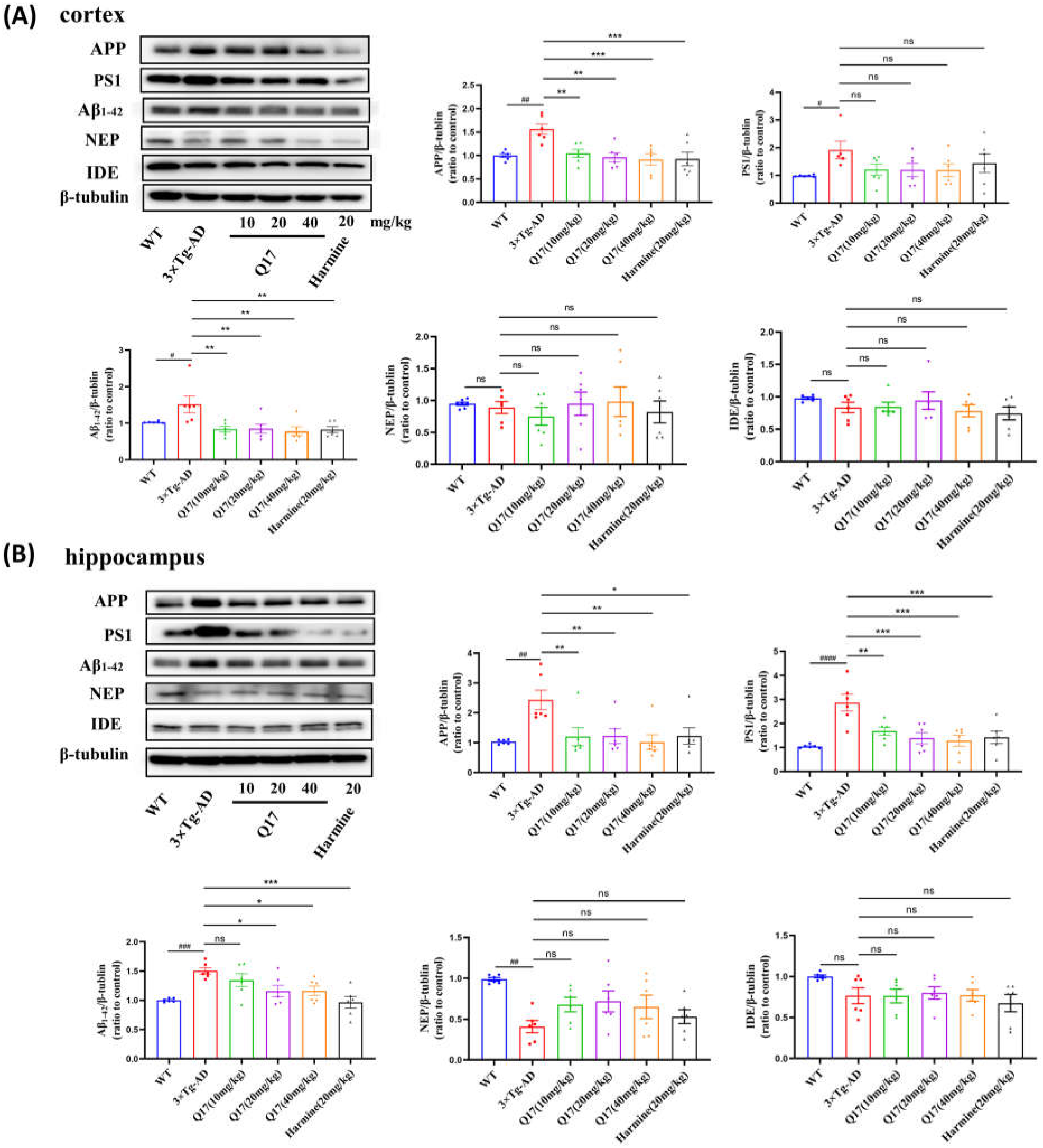
Compound **Q17** attenuated Aβ deposition in the hippocampus and cortex of 3×Tg-AD mice, shown by the western blot assay. The effect of **Q17** on Aβ protein deposition in the cortex (A) and hippocampus (B). Representative images are shown. Levels of APP expression, PS1 expression, Aβ_1-42_ expression, NEP expression, and IDE expression in the cortex and hippocampus were analyzed. The results are presented as mean ± SEM (n = 6). ^#^*p* < 0.05, ^##^*p* < 0.01, ^###^*p* < 0.001, ^####^*p*< 0.0001 vs WT group. \**p* < 0.05, \*\**p* < 0.01, \*\*\**p* < 0.001 vs 3×Tg-AD group. Harmine was used as a positive drug.

#### 3.3.5 Compound Q17 showed anti-AD efficacy by regulating DYRK1A and GSK-3β in 3×Tg-AD mice

As reported, tau phosphorylation by Dyrk1A is accompanied by further phosphorylation by GSK-3β [58]. We determined the expression of DYRK1A, GSK-3β, and GSK-3β (pSer9) by western blot as well. In 3×Tg-AD mice, we observed the expression of the above proteins increased in the cortex and hippocampus, but they were lowered down in a dose-dependent manner after the administration of compound **Q17** (Fig. 11A, B), suggesting that **Q17** can exert anti-AD effects in *vivo* by regulating DYRK1A and GSK-3β.

**Fig. 11.**
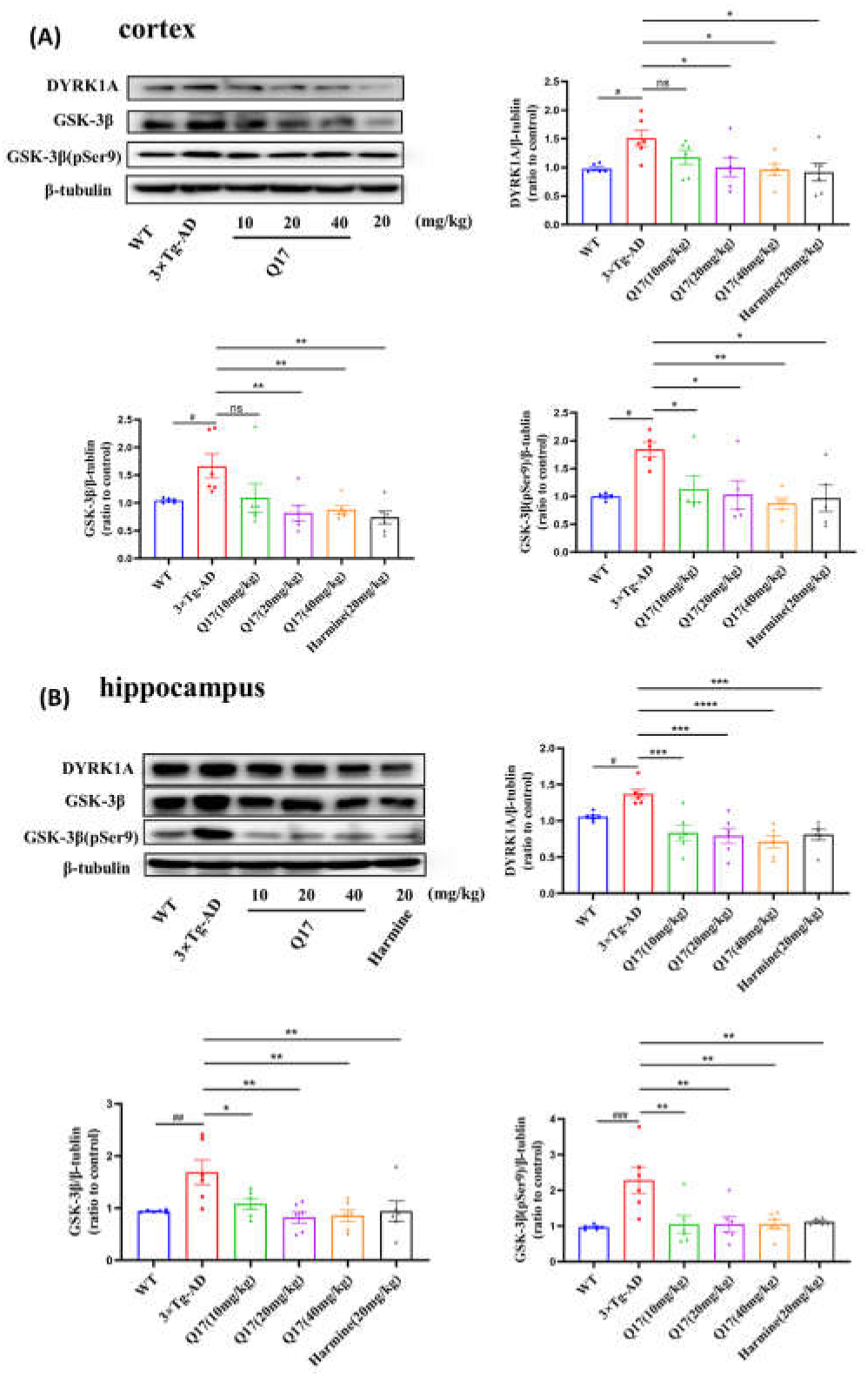
Compound **Q17** showed anti-AD effect by regulating the DYRK1A/GSK-3β pathway. The effect of **Q17** on the DYRK1A/GSK-3β pathway in the cortex (A) and hippocampus (B) of 3×Tg-AD mice. Representative bands are shown. Levels of DYRK1A expression, GSK-3β expression and GSK-3β (pSer9) expression in the cortex and hippocampus were analyzed. The results are presented as mean ± SEM (n = 6). ^#^*p* < 0.05, ^##^*p* < 0.01, ^####^*p*< 0.0001 vs WT group. \**p* < 0.05, \*\**p* < 0.01, \*\*\**p* < 0.001, \*\*\*\**p* < 0.0001 vs 3×Tg-AD group. Harmine was used as a positive drug.

## 4 Conclusion

AD has seriously threatened health of middle-aged and elderly people, bringing a significant economic burden to families and society [59]. However, research and development of anti-AD drugs is faced with great challenge. The only two anti-AD drugs approved by FDA, Donepezil and Memantine, can merely delay the progression of AD and is not available to cure that disease. Recently, DYRK1A is considered as a promising target for anti-AD drug discovery, as it phosphorylates tau protein and promotes Aβ generation [60, 61]. In addition, it has been observed that DYRK1A is overexpressed in the brains of AD patients [62], further confirming that DYRK1A is involved in AD pathogenesis. At present, several studies have suggested that DYRK1A inhibitors have great potential to treat AD [63–65]. However, the development of DYRK1A inhibitors for AD is still at the early stage. Currently, no DYRK1A inhibitor is approved as anti-AD therapy, only EGCG and SM07883 are in clinical trial [66, 67]. In our opinion, the lack of safe and diverse chemotypes hinders the rapid development of DYRK1A inhibitors as anti-AD drugs [68].

In this study, we carried out the high throughput VS, *in vitro* and *in vivo* biological evaluation, with the aim to fast identify novel-chemotype DYRK1A inhibitors that held the great potential to be developed into a new class of anti-AD drugs. To this end, we designed a rational workflow of high-throughput VS by (1) systematically analyzing 70 available co-crystal structures of DYRK1A for their ligand enrichment when they were coupled with FRED-based molecular docking and found out the structure with the PDB code of 7AJ2 was the most suitable; (2) generating a pharmacophore model and a shape model as filters, based on the selected co-crystal structure. Then, we applied the workflow to the *in silico* screening of the Specs chemical library and successfully discovered a DYRK1A inhibitor **Q17** (IC_50_:1.59 μM) whose chemotype had not been reported. By molecular dynamics simulation, this compound was predicted to form hydrogen bonds with GLU239, LEU241 and LYS188, which were consistent with the previous studies [69, 70] . The above data validated the value of the high throughput VS workflow in practice of new drug discovery.

Inspired by the novel chemotype, we were curious to see whether compound **Q17** was promising to treat AD. As a preliminary assay, we used OA-induced SH-SY5Y cells to test its neuroprotective activity and its related molecular mechanism. As expected, compound **Q17** significantly ameliorated OA-induced cell injury in SH-SY5Y cells, by inhibiting the expression of DYRK1A as well as the related proteins GSK-3β and pGSK-3β-ser9 thereby reducing Tau protein hyperphosphorylation. With 3×Tg-AD mice, we were able to evaluate *in vivo* efficacy of compound **Q17** for AD by the behavioral study (the NOR test, the Y-maze test, the MWM test and the PAT test), pathological examination and immunochemistry and western blot assays. Excitingly, it significantly improved cognitive dysfunction in 3×Tg-AD mice, ameliorated pathological changes, and reduced the expression of DYRK1A, GSK-3β and GSK-3β (pSer9), attenuated tau hyper-phosphorylation and Aβ deposition. Taken together, we have confirmed the identified compound **Q17** was effective at the animal level.

Though the bioactivity and efficacy of compound **Q17** were discovered for the first time, its chemical structure is known due to its source of the Specs chemical library. In addition, the DYRK1A inhibition is not particularly prominent compared with the other reported DYRK1A inhibitors, e.g. Harmine. To improve structural novelty and bioactivity, it is necessary to perform structural modification of compound **Q17**, which will be our future work.

In summary, we have discovered a DYRK1A inhibitor **Q17** by using computational modeling and biological evaluation. Compound **Q17** has been experimentally validated at the molecular level, cell level and the animal level as a novel class of DYRK1A inhibitors with promising anti-AD efficacy and thus worthy of further development.

## ASSOCIATED CONTENT AUTHOR INFORMATION

### Corresponding Author

*E-mail: J.X. and Xuehui Zhang or Hao Wang Address: Jie Xia, Institute of Materia Medica, Chinese Academy of Medical Sciences and Peking Union Medical College, Beijing 100050, China; Xuehui Zhang & Hao Wang. School of Pharmaceutical Sciences, Shandong First Medical University& Shandong Academy of Medical Sciences, Taian, Shandong 271016, China

## Acknowledgments

This work is supported by the CAMS Innovation Fund for Medical Sciences (Grant No. 2021-I2M-1-069) and National Natural Science Foundation of China (Grant No. 81603027 and 81973238).

## Author Contribution

Conception of the hypothesis: Jie Xia, Xuehui Zhang; Study supervision: Jie Xia, Xuehui Zhang, Hao Wang; Data acquisition, analysis and interpretation: Chenliang Qian, Yaling Wang, Hongwei Jin, Mingli Yao, Xinxin Si (for computational modeling), Chenliang Qian and Song Wu (for synthesis) and Nianzhuang Qiu, Tingting Guo, Mei Li, Tianyang Guo, Yuli Lv (for biology); Writing, review, and/or revision of the manuscript: Nianzhuang Qiu, Chenliang Qian, Jie Xia, Xuehui Zhang and Hao Wang. The manuscript is written through the contributions of all authors. All authors have given approval to the final version of the manuscript.

## Appendix A. Supplementary data

The supporting information to this article can be found online at https://doi.org/xxx

## Supporting Information

**Fig. S1.**
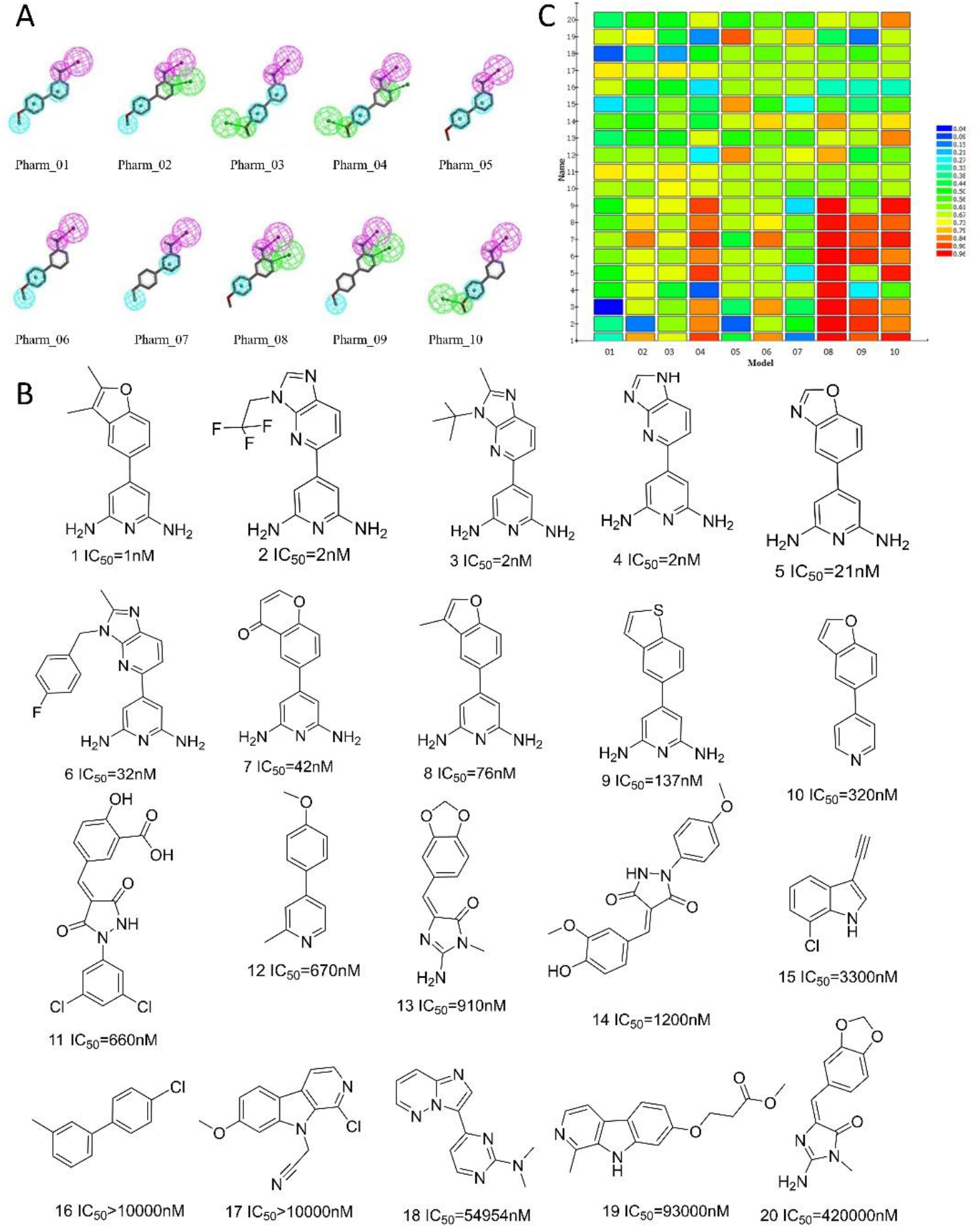
Pharmacophore models and performance evaluation. (A) 10 pharmacophore models constructed by Discovery Studio; (B) 20 compounds used to test pharmacophore models; (C) Heatmap to show model performance. It is easy to identify Pharm_08 as the optimal model as it assigned the maximal values to the active compounds and the minimal values to the inactive compounds

**Fig. S2.**
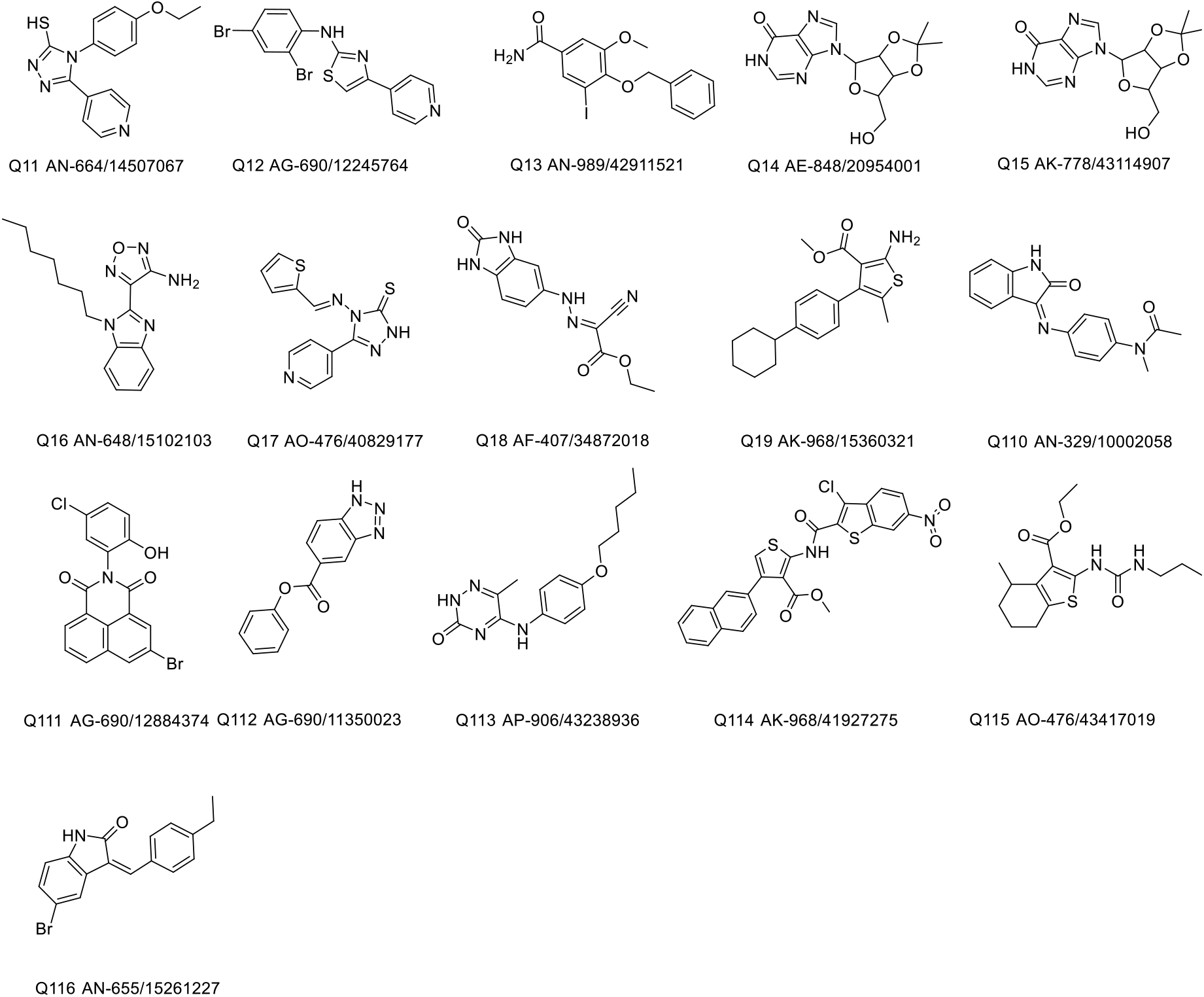
16 potential inhibitors suggested by virtual screening and available to purchase.

**Fig. S3.**
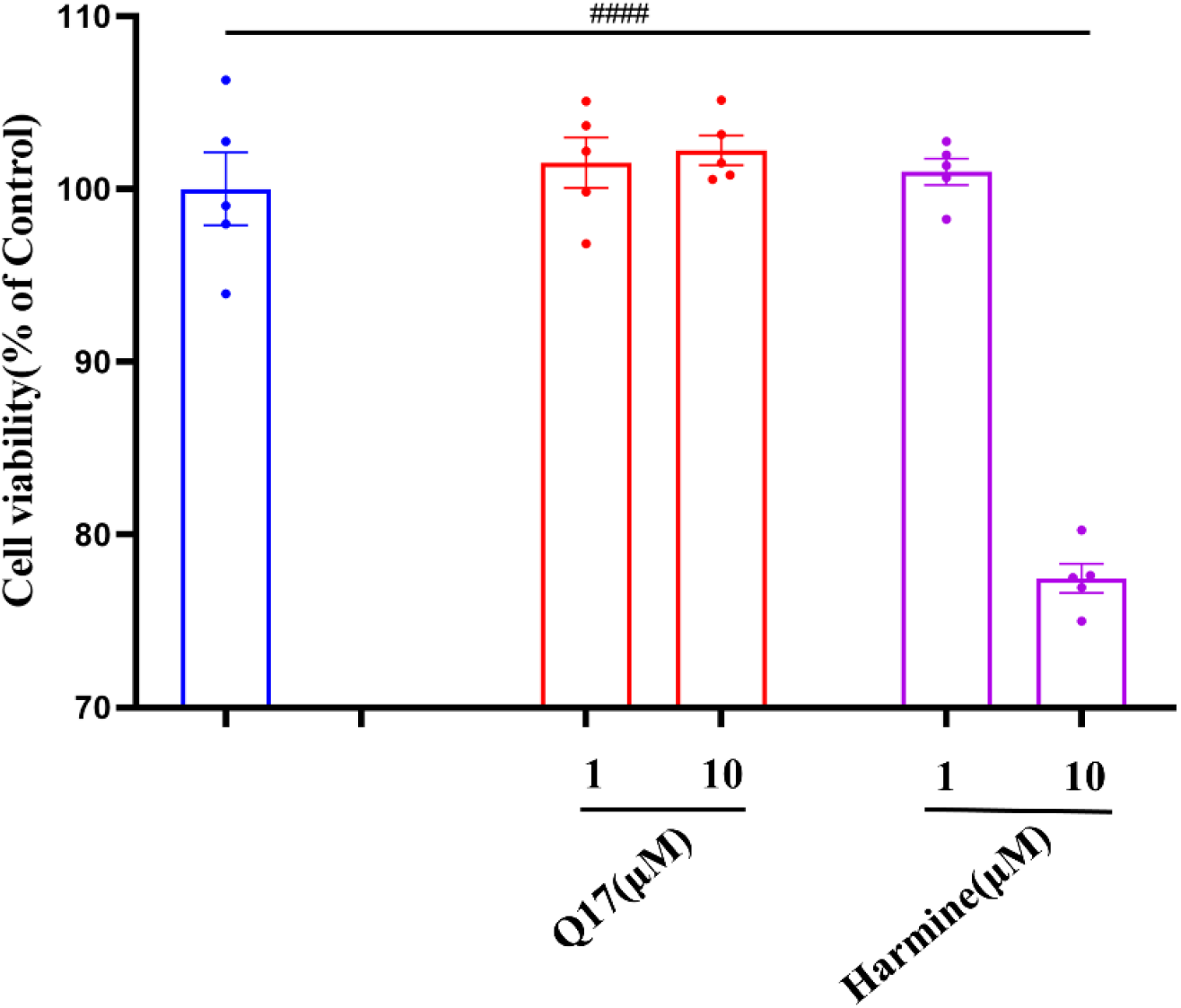
Effect of compound **Q17** itself on the viability of SH-SY5Y cells. The results are presented as mean ± SEM (n = 3). ** p < 0.01, **** p< 0.0001 vs control group. Harmine was used as the positive drug.

**Fig. S4.**
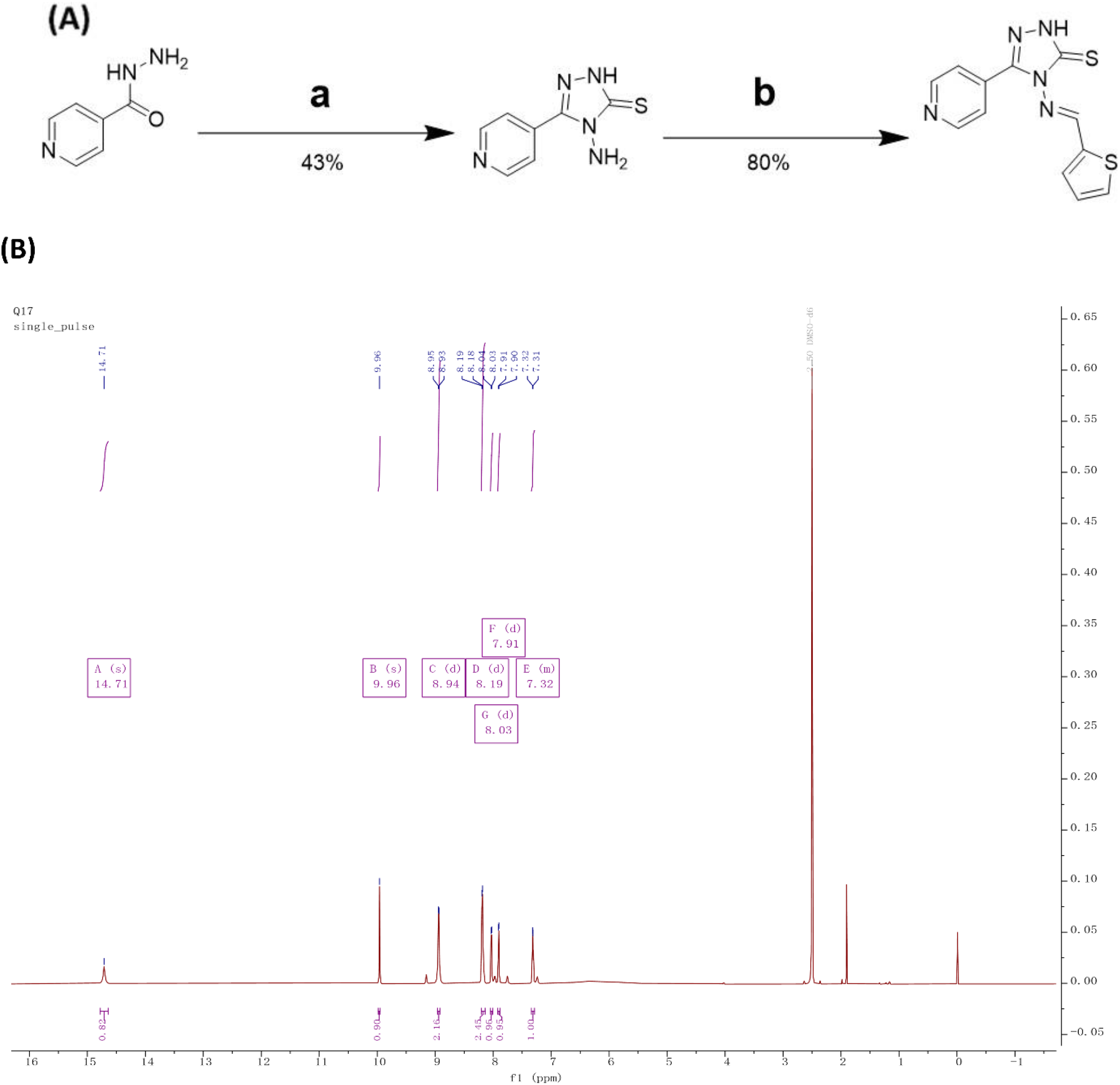
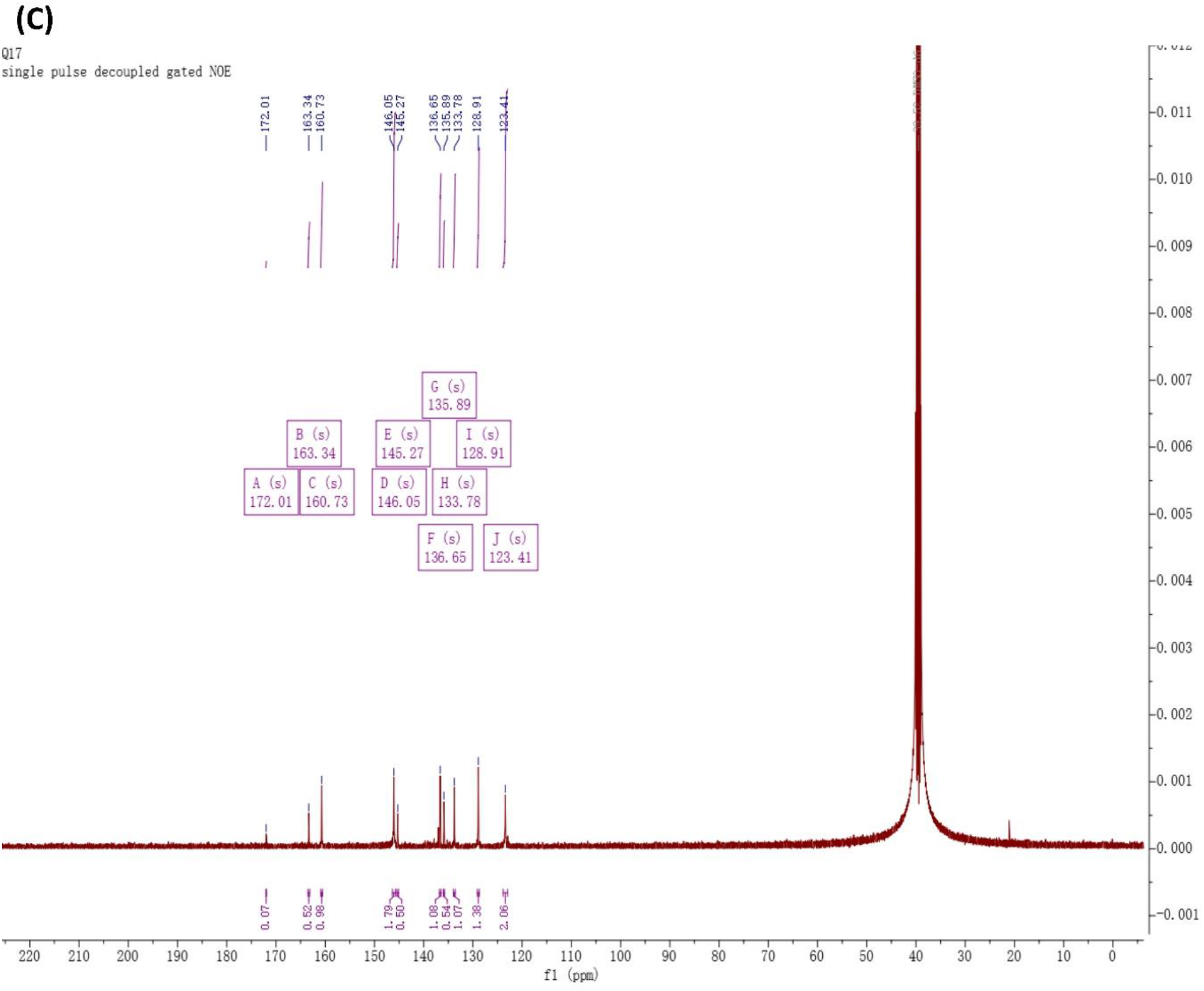
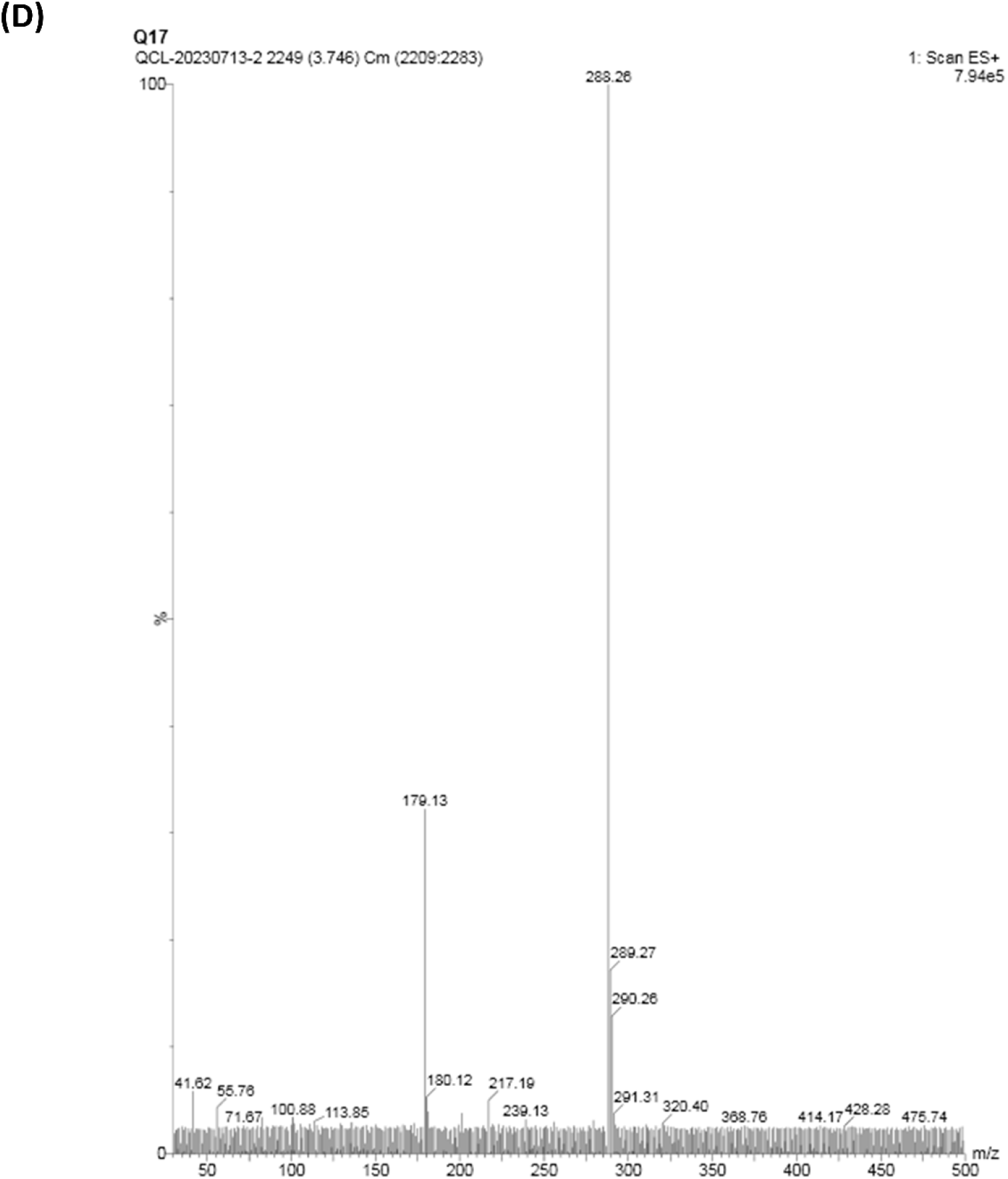
The synthetic route (A) and the structural validation (B-D) of compound Q17. Reagents and conditions: (a) CS2, KOH, MeOH, 0.5h; N_2_H_4_·H_2_O, H_2_O, reflux 4h; (b) CH_3_COOH; thiophene-2-carbaldehyde, reflux, 8h.^1^H NMR (500 MHz, DMSO) δ 14.71 (s, 1H), 9.96 (s, 1H), 8.94 (d, *J* = 5.6 Hz, 2H), 8.19 (d, *J* = 5.9 Hz, 2H), 8.03 (d, *J* = 5.0 Hz, 1H), 7.91 (d, *J* = 3.8 Hz, 1H), 7.34-7.29 (m, 1H). ^13^C NMR (126 MHz, DMSO) δ 172.01, 163.34, 160.73, 146.05, 145.27, 136.65, 135.89, 133.78, 128.91, 123.41. MS-ESI (m/z): [M + H] ^+^ calcd for C_12_H_9_N_5_S_2_, 288.03; found, 288.26.

**Table S1.**
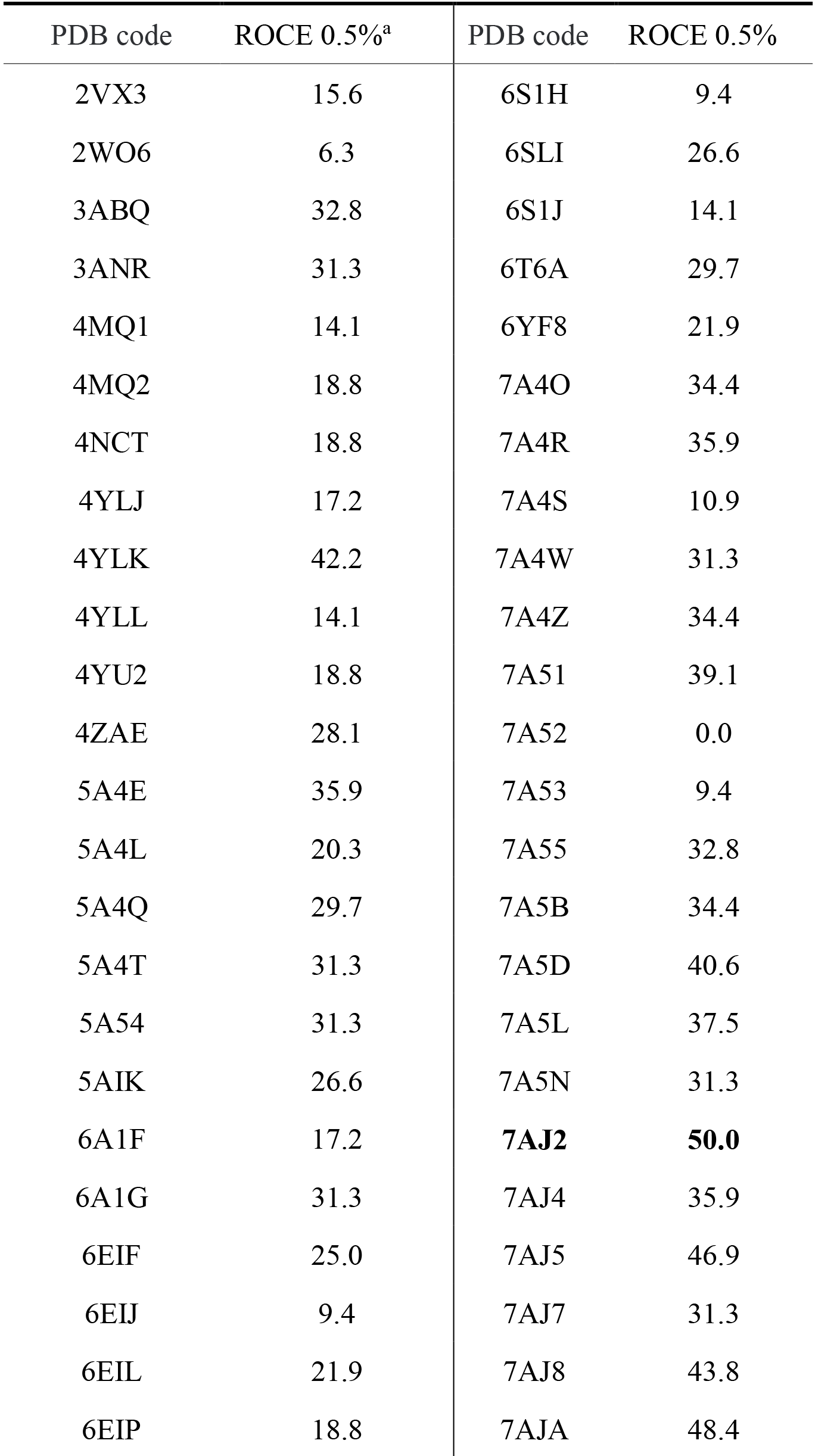
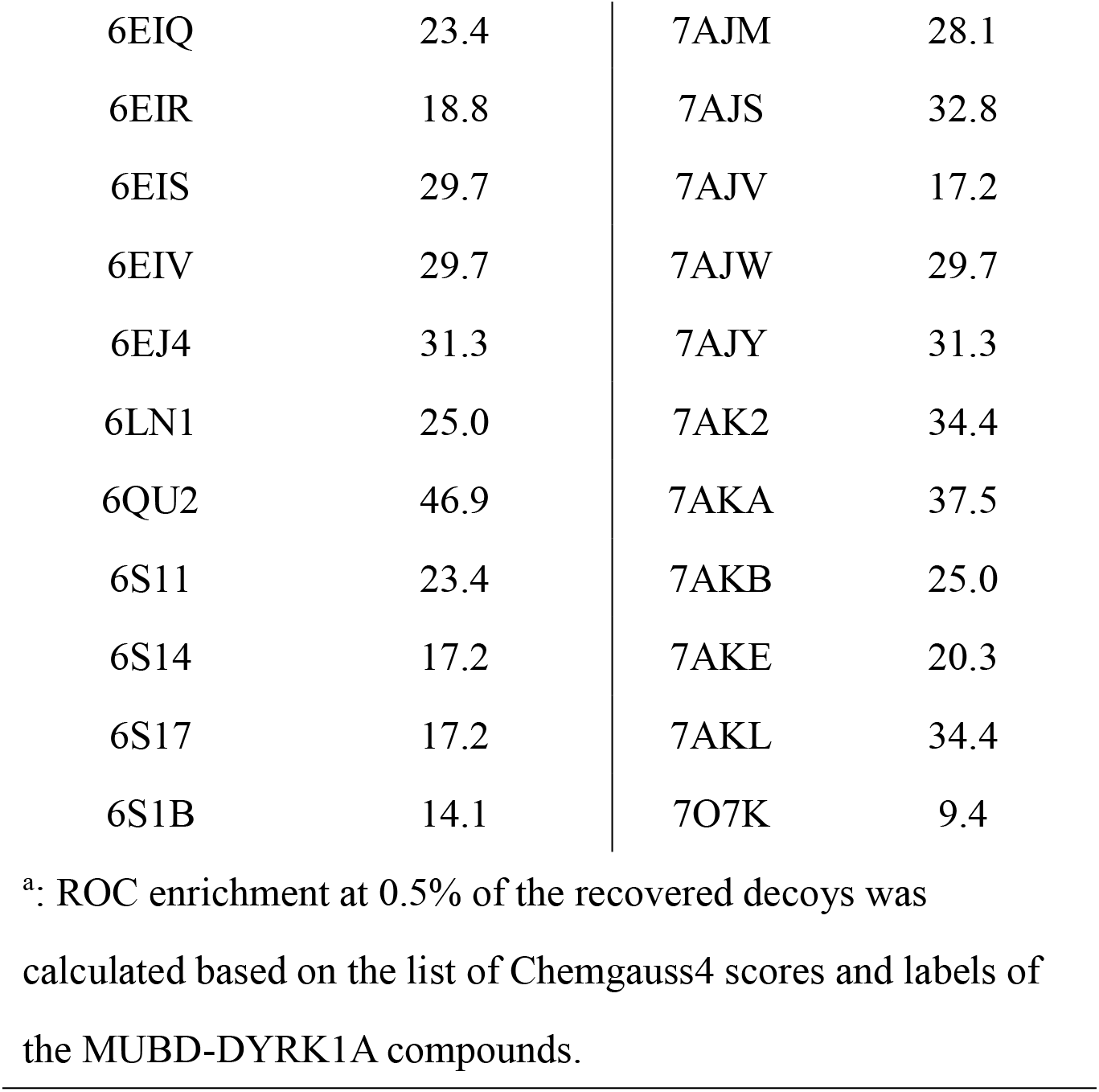
Ligand enrichments (ROCE0.5%) of FRED-based molecular docking approaches against 70 DYRK1A crystal structures.

**Table S2.** 16 purchased compounds: Specs ID and the FitValue, ShapeTanimoto (Rocs), Fred Chemgauss4 score.

|  | **SPECS ID** | **FitValue** | **ShapeTanimoto**<br>**(ROCS)** | **FRED Chemgauss4**<br>**score**<br>**(FRED)** |
| --- | --- | --- | --- | --- |
| **Q11** | AN-664/14507067 | 2.71477 | 0.651 | -12.9539 |
| **Q12** | AG-690/12245764 | 2.79341 | 0.736 | -12.1502 |
| **Q13** | AN-989/42911521 | 2.88491 | 0.678 | -12.8562 |
| **Q14** | AE-848/20954001 | 2.61947 | 0.755 | -13.8479 |
| **Q15** | AK-778/43114907 | 2.82797 | 0.673 | -12.2445 |
| **Q16** | AN-648/15102103 | 2.94995 | 0.688 | -12.5097 |
| **Q17** | AO-476/40829177 | 2.81998 | 0.654 | -14.6449 |
| **Q18** | AF-407/34872018 | 2.5007 | 0.78 | -12.3256 |
| **Q19** | AK-968/15360321 | 2.87747 | 0.655 | -12.2242 |
| **Q110** | AN-329/10002058 | 2.83581 | 0.674 | -12.2153 |
| **Q111** | AG-690/12884374 | 2.62526 | 0.698 | -12.7152 |
| **Q12** | AG-690/11350023 | 2.56372 | 0.819 | -13.1657 |
| **Q113** | AP-906/43238936 | 2.87524 | 0.754 | -11.9657 |
| **Q114** | AK-968/41927275 | 2.6796 | 0.842 | -11.9275 |
| **Q115** | AO-476/43417019 | 2.5254 | 0.661 | -12.4835 |
| **Q116** | AN-655/15261227 | 2.83155 | 0.656 | -13.3189 |

## References

[1] P. Scheltens, B. De Strooper, M. Kivipelto, H. Holstege, G. Chetelat, C.E. Teunissen, J. Cummings, W.M. van der Flier, Alzheimer’s disease, Lancet 397(10284) (2021) 1577–1590.

[2] J.A. Soria Lopez, H.M. Gonzalez, G.C. Leger, Alzheimer’s disease, Handb Clin Neurol 167 (2019) 231–255.

[3] S. Liu, M. Fan, Q. Zheng, S. Hao, L. Yang, Q. Xia, C. Qi, J. Ge, MicroRNAs in Alzheimer’s disease: Potential diagnostic markers and therapeutic targets, Biomed Pharmacother 148 (2022) 112681.

[4] P. Raina, P. Santaguida, A. Ismaila, C. Patterson, D. Cowan, M. Levine, L. Booker, M. Oremus, Effectiveness of cholinesterase inhibitors and memantine for treating dementia: evidence review for a clinical practice guideline, Ann Intern Med 148(5) (2008) 379–97.

[5] R. McShane, M.J. Westby, E. Roberts, N. Minakaran, L. Schneider, L.E. Farrimond, N. Maayan, J. Ware, J. Debarros, Memantine for dementia, Cochrane Database Syst Rev 3(3) (2019) CD003154.

[6] Y. Liu, L. Wang, F. Xie, X. Wang, Y. Hou, X. Wang, J. Liu, Overexpression of miR-26a-5p Suppresses Tau Phosphorylation and Abeta Accumulation in the Alzheimer’s Disease Mice by Targeting DYRK1A, Curr Neurovasc Res 17(3) (2020) 241–248.

[7] B. Melchior, G.K. Mittapalli, C. Lai, K. Duong-Polk, J. Stewart, B. Guner, B. Hofilena, A. Tjitro, S.D. Anderson, D.S. Herman, L. Dellamary, C.J. Swearingen, K.C. Sunil, Y. Yazici, Tau pathology reduction with SM07883, a novel, potent, and selective oral DYRK1A inhibitor: A potential therapeutic for Alzheimer’s disease, Aging Cell 18(5) (2019) e13000.

[8] D.R. Thal, J. Walter, T.C. Saido, M. Fandrich, Neuropathology and biochemistry of Abeta and its aggregates in Alzheimer’s disease, Acta Neuropathol 129(2) (2015) 167–82.

[9] I. Grundke-Iqbal, K. Iqbal, Y.C. Tung, M. Quinlan, H.M. Wisniewski, L.I. Binder, Abnormal phosphorylation of the microtubule-associated protein tau (tau) in Alzheimer cytoskeletal pathology, Proc Natl Acad Sci U S A 83(13) (1986) 4913–7.

[10] W.J. Song, L.R. Sternberg, C. Kasten-Sportes, M.L. Keuren, S.H. Chung, A.C. Slack, D.E. Miller, T.W. Glover, P.W. Chiang, L. Lou, D.M. Kurnit, Isolation of human and murine homologues of the Drosophila minibrain gene: human homologue maps to 21q22.2 in the Down syndrome "critical region", Genomics 38(3) (1996) 331–9.

[11] J. Wegiel, C.X. Gong, Y.W. Hwang, The role of DYRK1A in neurodegenerative diseases, FEBS J 278(2) (2011) 236–45.

[12] G.V. Johnson, J.A. Hartigan, Tau protein in normal and Alzheimer’s disease brain: an update, J Alzheimers Dis 1(4-5) (1999) 329–51.

[13] T. Liu, Y. Wang, J. Wang, C. Ren, H. Chen, J. Zhang, DYRK1A inhibitors for disease therapy: Current status and perspectives, Eur J Med Chem 229 (2022) 114062.

[14] S.R. Ryoo, H.J. Cho, H.W. Lee, H.K. Jeong, C. Radnaabazar, Y.S. Kim, M.J. Kim, M.Y. Son, H. Seo, S.H. Chung, W.J. Song, Dual-specificity tyrosine(Y)-phosphorylation regulated kinase 1A-mediated phosphorylation of amyloid precursor protein: evidence for a functional link between Down syndrome and Alzheimer’s disease, J Neurochem 104(5) (2008) 1333–44.

[15] M. Rammohan, E. Harris, R.S. Bhansali, E. Zhao, L.S. Li, J.D. Crispino, The chromosome 21 kinase DYRK1A: emerging roles in cancer biology and potential as a therapeutic target, Oncogene 41(14) (2022) 2003–2011.

[16] A. Walte, K. Ruben, R. Birner-Gruenberger, C. Preisinger, S. Bamberg-Lemper, N. Hilz, F. Bracher, W. Becker, Mechanism of dual specificity kinase activity of DYRK1A, FEBS J 280(18) (2013) 4495–511.

[17] J. Wegiel, K. Dowjat, W. Kaczmarski, I. Kuchna, K. Nowicki, J. Frackowiak, B. Mazur Kolecka, J. Wegiel, W.P. Silverman, B. Reisberg, M. Deleon, T. Wisniewski, C.X. Gong, F. Liu, T. Adayev, M.C. Chen-Hwang, Y.W. Hwang, The role of overexpressed DYRK1A protein in the early onset of neurofibrillary degeneration in Down syndrome, Acta Neuropathol 116(4) (2008) 391–407.

[18] Y.S. Ryu, S.Y. Park, M.S. Jung, S.H. Yoon, M.Y. Kwen, S.Y. Lee, S.H. Choi, C. Radnaabazar, M.K. Kim, H. Kim, K. Kim, W.J. Song, S.H. Chung, Dyrk1A-mediated phosphorylation of Presenilin 1: a functional link between Down syndrome and Alzheimer’s disease, J Neurochem 115(3) (2010) 574–84.

[19] J. Park, K.C. Chung, New Perspectives of Dyrk1A Role in Neurogenesis and Neuropathologic Features of Down Syndrome, Exp Neurobiol 22(4) (2013) 244–8.

[20] J. Park, W.J. Song, K.C. Chung, Function and regulation of Dyrk1A: towards understanding Down syndrome, Cell Mol Life Sci 66(20) (2009) 3235–40.

[21] S.A. Almatroodi, A. Almatroudi, A.A. Khan, F.A. Alhumaydhi, M.A. Alsahli, A.H. Rahmani, Potential Therapeutic Targets of Epigallocatechin Gallate (EGCG), the Most Abundant Catechin in Green Tea, and Its Role in the Therapy of Various Types of Cancer, Molecules 25(14) (2020).

[22] S. Zhang, Q. Zhu, J.Y. Chen, D. OuYang, J.H. Lu, The pharmacological activity of epigallocatechin-3-gallate (EGCG) on Alzheimer’s disease animal model: A systematic review, Phytomedicine 79 (2020) 153316.

[23] S. Stotani, F. Giordanetto, F. Medda, DYRK1A inhibition as potential treatment for Alzheimer’s disease, Future Med Chem 8(6) (2016) 681–96.

[24] S. Li, Y. Zhang, G. Deng, Y. Wang, S. Qi, X. Cheng, Y. Ma, Y. Xie, C. Wang, Exposure Characteristics of the Analogous beta-Carboline Alkaloids Harmaline and Harmine Based on the Efflux Transporter of Multidrug Resistance Protein 2, Front Pharmacol 8 (2017) 541.

[25] A. Wurzlbauer, K. Ruben, E. Gurdal, A. Chaikuad, S. Knapp, W. Sippl, W. Becker, F. Bracher, How to Separate Kinase Inhibition from Undesired Monoamine Oxidase A Inhibition-The Development of the DYRK1A Inhibitor AnnH75 from the Alkaloid Harmine, Molecules 25(24) (2020).

[26] M.M. Airaksinen, A. Lecklin, V. Saano, L. Tuomisto, J. Gynther, Tremorigenic effect and inhibition of tryptamine and serotonin receptor binding by beta-carbolines, Pharmacol Toxicol 60(1) (1987) 5–8.

[27] J. Xia, E.L. Tilahun, T.E. Reid, L. Zhang, X.S. Wang, Benchmarking methods and data sets for ligand enrichment assessment in virtual screening, Methods 71 (2015) 146–57.

[28] D. Wang, F. Wang, Y. Tan, L. Dong, L. Chen, W. Zhu, H. Wang, Discovery of potent small molecule inhibitors of DYRK1A by structure-based virtual screening and bioassay, Bioorg Med Chem Lett 22(1) (2012) 168–71.

[29] T. Koyama, N. Yamaotsu, I. Nakagome, S.I. Ozawa, T. Yoshida, D. Hayakawa, S. Hirono, Multi-step virtual screening to develop selective DYRK1A inhibitors, J Mol Graph Model 72 (2017) 229–239.

[30] M. Choi, A.K. Kim, Y. Ham, J.Y. Lee, D. Kim, A. Yang, M.J. Jo, E. Yoon, J.N. Heo, S.B. Han, M.H. Ki, K.S. Lee, S. Cho, Aristolactam BIII, a naturally derived DYRK1A inhibitor, rescues Down syndrome-related phenotypes, Phytomedicine 92 (2021) 153695.

[31] J. Xia, Z. Wang, Y. Huan, W. Xue, X. Wang, Y. Wang, Z. Liu, J.H. Hsieh, L. Zhang, S. Wu, Z. Shen, H. Zhang, X.S. Wang, Pose Filter-Based Ensemble Learning Enables Discovery of Orally Active, Nonsteroidal Farnesoid X Receptor Agonists, J Chem Inf Model 60(3) (2020) 1202–1214.

[32] T. Qin, X. Gao, L. Lei, J. Feng, W. Zhang, Y. Hu, Z. Shen, Z. Liu, Y. Huan, S. Wu, J. Xia, L. Zhang, Machine learning- and structure-based discovery of a novel chemotype as FXR agonists for potential treatment of nonalcoholic fatty liver disease, Eur J Med Chem 252 (2023) 115307.

[33] J. Xia, S. Li, Y. Ding, S. Wu, X.S. Wang, MUBD-DecoyMaker 2.0: A Python GUI Application to Generate Maximal Unbiased Benchmarking Data Sets for Virtual Drug Screening, Mol Inform 39(4) (2020) e1900151.

[34] J. Xia, E.L. Tilahun, E.H. Kebede, T.E. Reid, L. Zhang, X.S. Wang, Comparative modeling and benchmarking data sets for human histone deacetylases and sirtuin families, J Chem Inf Model 55(2) (2015) 374–88.

[35] P.C. Hawkins, A.G. Skillman, G.L. Warren, B.A. Ellingson, M.T. Stahl, Conformer generation with OMEGA: algorithm and validation using high quality structures from the Protein Databank and Cambridge Structural Database, J Chem Inf Model 50(4) (2010) 572–84.

[36] M. McGann, FRED pose prediction and virtual screening accuracy, J Chem Inf Model 51(3) (2011) 578–96.

[37] J. Xia, J.H. Hsieh, H. Hu, S. Wu, X.S. Wang, The Development of Target-Specific Pose Filter Ensembles To Boost Ligand Enrichment for Structure-Based Virtual Screening, J Chem Inf Model 57(6) (2017) 1414–1425.

[38] M.A. DeTure, D.W. Dickson, The neuropathological diagnosis of Alzheimer’s disease, Mol Neurodegener 14(1) (2019) 32.

[39] P.C. Hawkins, A.G. Skillman, A. Nicholls, Comparison of shape-matching and docking as virtual screening tools, J Med Chem 50(1) (2007) 74–82.

[40] J. Venhorst, S. Nunez, J.W. Terpstra, C.G. Kruse, Assessment of scaffold hopping efficiency by use of molecular interaction fingerprints, J Med Chem 51(11) (2008) 3222–9.

[41] T. Qin, X. Gao, L. Lei, W. Zhang, J. Feng, X. Wang, Z. Shen, Z. Liu, Y. Huan, S. Wu, J. Xia, L. Zhang, Structural optimization and biological evaluation of 1-adamantylcarbonyl-4-phenylpiperazine derivatives as FXR agonists for NAFLD, Eur J Med Chem 245(Pt 1) (2023) 114903.

[42] M. Christen, P.H. Hunenberger, D. Bakowies, R. Baron, R. Burgi, D.P. Geerke, T.N. Heinz, M.A. Kastenholz, V. Krautler, C. Oostenbrink, C. Peter, D. Trzesniak, W.F. van Gunsteren, The GROMOS software for biomolecular simulation: GROMOS05, J Comput Chem 26(16) (2005) 1719–51.

[43] L.S. Dodda, I. Cabeza de Vaca, J. Tirado-Rives, W.L. Jorgensen, LigParGen web server: an automatic OPLS-AA parameter generator for organic ligands, Nucleic Acids Res 45(W1) (2017) W331–W336.

[44] W.R.P. Scott, P.H. Hunenberger, I.G. Tironi, A.E. Mark, S.R. Billeter, J. Fennen, A.E. Torda, T. Huber, P. Kruger, W.F. van Gunsteren, The GROMOS biomolecular simulation program package, J. Phys. Chem. A 103(19) (1999) 3596–3607.

[45] S.B. Zhu, S. Yao, J.B. Zhu, S. Singh, G.W. Robinson, A flexible/polarizable simple point charge water model, J Phys Chem 95(16) (1991) 6211–6217.

[46] M.A. Hussein, R.M. Shaker, M.A. Ameen, M.F. Mohammed, Synthesis, anti-inflammatory, analgesic, and antibacterial activities of some triazole, triazolothiadiazole, and triazolothiadiazine derivatives, Arch Pharm Res 34(8) (2011) 1239–50.

[47] C. Lechner, M. Flasshoff, H. Falke, L. Preu, N. Loaec, L. Meijer, S. Knapp, A. Chaikuad, C. Kunick, [b]-Annulated Halogen-Substituted Indoles as Potential DYRK1A Inhibitors, Molecules 24(22) (2019).

[48] R.A. Bevins, J. Besheer, Object recognition in rats and mice: a one-trial non-matching-to-sample learning task to study ’recognition memory’, Nat Protoc 1(3) (2006) 1306–11.

[49] J.C. Carroll, E.R. Rosario, L. Chang, F.Z. Stanczyk, S. Oddo, F.M. LaFerla, C.J. Pike, Progesterone and estrogen regulate Alzheimer-like neuropathology in female 3xTg-AD mice, J Neurosci 27(48) (2007) 13357–65.

[50] X. Dou, H. Huang, Y. Li, L. Jiang, Y. Wang, H. Jin, N. Jiao, L. Zhang, L. Zhang, Z. Liu, Multistage Screening Reveals 3-Substituted Indolin-2-one Derivatives as Novel and Isoform-Selective c-Jun N-terminal Kinase 3 (JNK3) Inhibitors: Implications to Drug Discovery for Potential Treatment of Neurodegenerative Diseases, J Med Chem 62(14) (2019) 6645–6664.

[51] X. Jin, M.Y. Liu, D.F. Zhang, X. Zhong, K. Du, P. Qian, W.F. Yao, H. Gao, M.J. Wei, Baicalin mitigates cognitive impairment and protects neurons from microglia-mediated neuroinflammation via suppressing NLRP3 inflammasomes and TLR4/NF-kappaB signaling pathway, CNS Neurosci Ther 25(5) (2019) 575–590.

[52] Y.P. Wang, X.T. Li, S.J. Liu, X.W. Zhou, X.C. Wang, J.Z. Wang, Melatonin ameliorated okadaic-acid induced Alzheimer-like lesions, Acta Pharmacol. Sin. 25(3) (2004) 276–280.

[53] D. He, H. Wu, Y. Wei, W. Liu, F. Huang, H. Shi, B. Zhang, X. Wu, C. Wang, Effects of harmine, an acetylcholinesterase inhibitor, on spatial learning and memory of APP/PS1 transgenic mice and scopolamine-induced memory impairment mice, Eur J Pharmacol 768 (2015) 96–107.

[54] J.T. Yang, Z.J. Wang, H.Y. Cai, L. Yuan, M.M. Hu, M.N. Wu, J.S. Qi, Sex Differences in Neuropathology and Cognitive Behavior in APP/PS1/tau Triple-Transgenic Mouse Model of Alzheimer’s Disease, Neurosci. Bull. 34(5) (2018) 736–746.

[55] Y. Zhao, R. Qian, J. Zhang, F. Liu, K. Iqbal, C.L. Dai, C.X. Gong, Young blood plasma reduces Alzheimer’s disease-like brain pathologies and ameliorates cognitive impairment in 3xTg-AD mice, Alzheimers Res. Ther. 12(1) (2020) 13.

[56] M.F. Falangola, X.J. Nie, R. Ward, E.T. McKinnon, S. Dhiman, P.J. Nietert, J.A. Helpern, J.H. Jensen, Diffusion MRI detects early brain microstructure abnormalities in 2-month-old 3xTg-AD mice, NMR Biomed. 33(9) (2020) 12.

[57] Y.H. Liu, X.L. Bu, C.R. Liang, Y.R. Wang, T. Zhang, S.S. Jiao, F. Zeng, X.Q. Yao, H.D. Zhou, J. Deng, Y.J. Wang, An N-terminal antibody promotes the transformation of amyloid fibrils into oligomers and enhances the neurotoxicity of amyloid-beta: the dust-raising effect, J. Neuroinflamm. 12 (2015) 8.

[58] F. Liu, Z.H. Liang, J. Wegiel, Y.W. Hwang, K. Iqbal, I. Grundke-Iqbal, N. Ramakrishna, C.X. Gong, Overexpression of Dyrk1A contributes to neurofibrillary degeneration in Down syndrome, Faseb J. 22(9) (2008) 3224–3233.

[59] A.A. Tahami Monfared, M.J. Byrnes, L.A. White, Q. Zhang, The Humanistic and Economic Burden of Alzheimer’s Disease, Neurol Ther 11(2) (2022) 525–551.

[60] R. Kimura, K. Kamino, M. Yamamoto, A. Nuripa, T. Kida, H. Kazui, R. Hashimoto, T. Tanaka, T. Kudo, H. Yamagata, Y. Tabara, T. Miki, H. Akatsu, K. Kosaka, E. Funakoshi, K. Nishitomi, G. Sakaguchi, A. Kato, H. Hattori, T. Uema, M. Takeda, The DYRK1A gene, encoded in chromosome 21 Down syndrome critical region, bridges between beta-amyloid production and tau phosphorylation in Alzheimer disease, Hum. Mol. Genet. 16(1) (2007) 15–23.

[61] Y.L. Woods, P. Cohen, W. Becker, R. Jakes, M. Goedert, X.M. Wang, C.G. Proud, The kinase DYRK phosphorylates protein-synthesis initiation factor eIF2B epsilon at Ser(539) and the microtubule-associated protein tau at Thr(212): potential role for DYRK as a glycogen synthase kinase 3-priming kinase, Biochem. J. 355 (2001) 609–615.

[62] M. Moreau, M. Carmona-Iragui, M. Altuna, L. Dalzon, I. Barroeta, M. Vilaire, S. Durand, J. Fortea, A.S. Rebillat, N. Janel, DYRK1A and Activity-Dependent Neuroprotective Protein Comparative Diagnosis Interest in Cerebrospinal Fluid and Plasma in the Context of Alzheimer-Related Cognitive Impairment in Down Syndrome Patients, Biomedicines 10(6) (2022).

[63] X. Liu, L.Y. Lai, J.X. Chen, X. Li, N. Wang, L.J. Zhou, X.W. Jiang, X.L. Hu, W.W. Liu, X.M. Jiao, Z.T. Qi, W.J. Liu, L.M. Wu, Y.G. Huang, Z.H. Xu, Q.C. Zhao, An inhibitor with GSK3beta and DYRK1A dual inhibitory properties reduces Tau hyperphosphorylation and ameliorates disease in models of Alzheimer’s disease, Neuropharmacology 232 (2023) 109525.

[64] H.J. Lee, H. Woo, H.E. Lee, H. Jeon, K.Y. Ryu, J.H. Nam, S.G. Jeon, H. Park, J.S. Lee, K.M. Han, S.M. Lee, J. Kim, R.J. Kang, Y.H. Lee, J.I. Kim, H.S. Hoe, The novel DYRK1A inhibitor KVN93 regulates cognitive function, amyloid-beta pathology, and neuroinflammation, Free Radic Biol Med 160 (2020) 575–595.

[65] M.M. de Souza, A.R. Cenci, K.F. Teixeira, V. Machado, M.C.G. Mendes Schuler, A.E. Goncalves, A. Paula Dalmagro, C. Andre Cazarin, L.L. Gomes Ferreira, A.S. de Oliveira, A.D. Andricopulo, DYRK1A Inhibitors and Perspectives for the Treatment of Alzheimer’s Disease, Curr Med Chem 30(6) (2023) 669–688.

[66] C. Secker, A.Y. Motzny, S. Kostova, A. Buntru, L. Helmecke, L. Reus, R. Steinfort, L. Brusendorf, A. Boeddrich, N. Neuendorf, L. Diez, P. Schmieder, A. Schulz, C. Czekelius, E.E. Wanker, The polyphenol EGCG directly targets intracellular amyloid-beta aggregates and promotes their lysosomal degradation, J. Neurochem. 166(2) (2023) 294–317.

[67] B. Melchior, G.K. Mittapalli, C. Lai, K. Duong-Polk, J. Stewart, B. Guner, B. Hofilena, A. Tjitro, S.D. Anderson, D.S. Herman, L. Dellamary, C.J. Swearingen, K.C. Sunil, Y. Yazici, Tau pathology reduction with SM07883, a novel, potent, and selective oral DYRK1A inhibitor: A potential therapeutic for Alzheimer’s disease, Aging Cell 18(5) (2019) 14.

[68] S. Stotani, F. Giordanetto, F. Medda, DYRK1A inhibition as potential treatment for Alzheimer’s disease, Future Med. Chem. 8(6) (2016) 681–696.

[69] Y. Ogawa, Y. Nonaka, T. Goto, E. Ohnishi, T. Hiramatsu, I. Kii, M. Yoshida, T. Ikura, H. Onogi, H. Shibuya, T. Hosoya, N. Ito, M. Hagiwara, Development of a novel selective inhibitor of the Down syndrome-related kinase Dyrk1A, Nat Commun 1 (2010) 86.

[70] M. Soundararajan, A.K. Roos, P. Savitsky, P. Filippakopoulos, A.N. Kettenbach, J.V. Olsen, S.A. Gerber, J. Eswaran, S. Knapp, J.M. Elkins, Structures of Down syndrome kinases, DYRKs, reveal mechanisms of kinase activation and substrate recognition, Structure 21(6) (2013) 986–96.

